# Themis dominates T cell exhaustion by regulation of TCR and PD-1 signaling

**DOI:** 10.1101/2025.02.17.634430

**Authors:** Jian Tang, Xian Jia, Taoling Zeng, Yuzhou Bao, Yanyan Hu, Junchen Dong, Xiaozheng Xu, Xiao Lei Chen, Yongchao Liu, Jiayu Wang, Bowen Hou, Yuzhen Zhu, Guo-Feng Fu, Qifan Zheng, Yongling Chen, Wanyun Li, Tong Ren, Lei Zhang, Shan Jiang, Yu Cong, Minxue Quan, Chaonan Yan, Shuo Lin, Ning Wang, Siyi Liu, Chenghao Huang, Lichao Hou, John Teijaro, Joanna Brzostek, Namrata Gautam, Changchun Xiao, Yan Wang, Gang Li, Bing Xu, Enfu Hui, Hong-Rui Wang, Nicholas R. J. Gascoigne, Guo Fu

## Abstract

T cell exhaustion is important to protect the host from immunopathology during chronic viral infection, but it also impairs T cell anti-tumor immunity^1–5^. A fundamental unresolved question is whether and how T cell exhaustion is determined at the onset of TCR signaling^6–8^. Here we report an unexpected role of Themis, a TCR-proximal signaling molecule^9^, in T cell exhaustion. Chronic viral infection in mice usually leads to T cell exhaustion and survival of the host. Surprisingly, Themis T-cell conditional knockout mice died from severe CD8^+^-dependent lung immunopathology in chronic viral infection, showing Themis’ importance in establishing T cell exhaustion. We found that Themis-deficient CD8^+^ T cells were hyperactivated at the single-cell level - producing more TNF and IFNγ compared to wild-type counterparts - but defective in population-level expansion. Moreover, TCF-1 and TOX expression were inhibited in Themis-deficient CD8^+^ T cells, thereby impairing differentiation of exhausted T cell precursors (T-pex) and maintenance of terminally exhausted T cells (T-ex), respectively. Mechanistically, Themis initially promotes TCR signaling to induce PD-1 expression and subsequently mediates PD-1 signaling. In the latter, Themis binds to PD-1 and promotes PD-1 phosphorylation and its recruitment of SHP2, thereby acting as a negative regulator to inhibit T cell effector functions. Without Themis, the orderly regulation of TCR and PD-1 signaling, and therefore exhaustion, is disrupted. Thus, our results unequivocally demonstrate that Themis-mediated early TCR signaling plays a decisive role in T cell exhaustion and provide a novel mechanism of PD-1 signaling through Themis.

---

T cell exhaustion is a state in which T cells become functionally impaired due to prolonged exposure to antigens^1,2^. The main features of T cell exhaustion include the loss of effector cytokine production and the expression of multiple co-inhibitory molecules^2,3^. Despite tremendous progress in the study of T cell exhaustion^2,4,5^, a fundamental unresolved question is how T cell exhaustion begins. Recent studies suggest that T cell exhaustion may initiate earlier than previously expected in chronic viral infections^6^ and tumors^7^, and may involve early TCR signaling^8^. Themis is a TCR proximal signaling molecule required for thymocyte development^9–14^, partly mediated through its interaction with the phosphatase SHP1^14–17^. Themis deficiency also results in severe defects in CD8^+^ T cell homeostasis^18^, cytokine responsiveness^19^, and signaling^20^. Despite these defects, Themis-deficient CD8^+^ T cells still mount normal antiviral immune responses to acute LCMV infection and even surprisingly produce more effector cytokines^21^. Moreover, we found that Themis binds to PD-1, a hallmark molecule of T cell exhaustion^22^, in thymocytes^21^, prompting us to consider whether Themis is involved in T cell exhaustion and PD-1 signaling. Here we reported that Themis T-cell conditional knockout mice succumb to chronic LCMV C13 infection due to CD8^+^ T cell mediated lung damage. Themis-deficient CD8^+^ T cells exhibited enhanced cytokine production but impaired clonal expansion, rendering these cells incapable of clearing the virus but sufficient to induce mortality. Mechanistically, Themis initially promotes TCR signaling to induce PD-1 expression and subsequently mediates PD-1 phosphorylation and signaling. Without Themis, the orderly regulation of TCR and PD-1 signaling is disrupted, as is T cell exhaustion.

## Results

### Themis T-cell conditional knockout mice succumb to chronic LCMV infection due to CD8^+^ T cell-mediated pathology

To investigate whether Themis is involved in T cell exhaustion, we infected Themis T-cell conditional knockout (cKO) mice^18^ or their control (WT) mice with LCMV C13. We found that while almost all WT mice survived infection, approximately 79% of cKO mice died 8 to 10 days post infection (dpi) (Fig. 1a). Autopsy revealed elevated levels of multiple proinflammatory cytokines in bronchoalveolar lavage fluid (Extended Data Fig. 1a), but comparable virus titers (Extended Data Fig. 1b). To address this question, we performed imaging mass cytometry^23^ on lung sections from cKO mice (Extended Data Fig. 1c and d). We found that the bronchi and blood vessels were mostly intact and identifiable in WT mice, but not in cKO mice (Fig. 1b). Strikingly, in the lung of cKO mice, we observed intense TUNEL signals that colocalized primarily with E-Cadherin and granzyme B, whereas the adjacent αSMA staining disappeared (Fig. 1b, panel I; Extended Data Fig. 1e, panel I), indicating severe cell death. The pulmonary vessels of cKO mice lacked TUNEL signals but were thinner than those of WT mice (Extended Data Fig. 1e, panel II). Neutrophil and monocyte infiltration was mild in both mouse strains (Extended Data Fig. 1e, panel III), whereas CD8^+^ T cell infiltration was prominent and CD4^+^ T cell infiltration was only seen in cKO mice (Fig. 1b, panel II). PD-L1 expression pattern was comparable (Extended Data Fig. 1e, panel IV). Confocal microscopy confirmed the exclusive staining of TUNEL in the lung of cKO mice (Extended Data Fig. 1f).

**Fig. 1.**
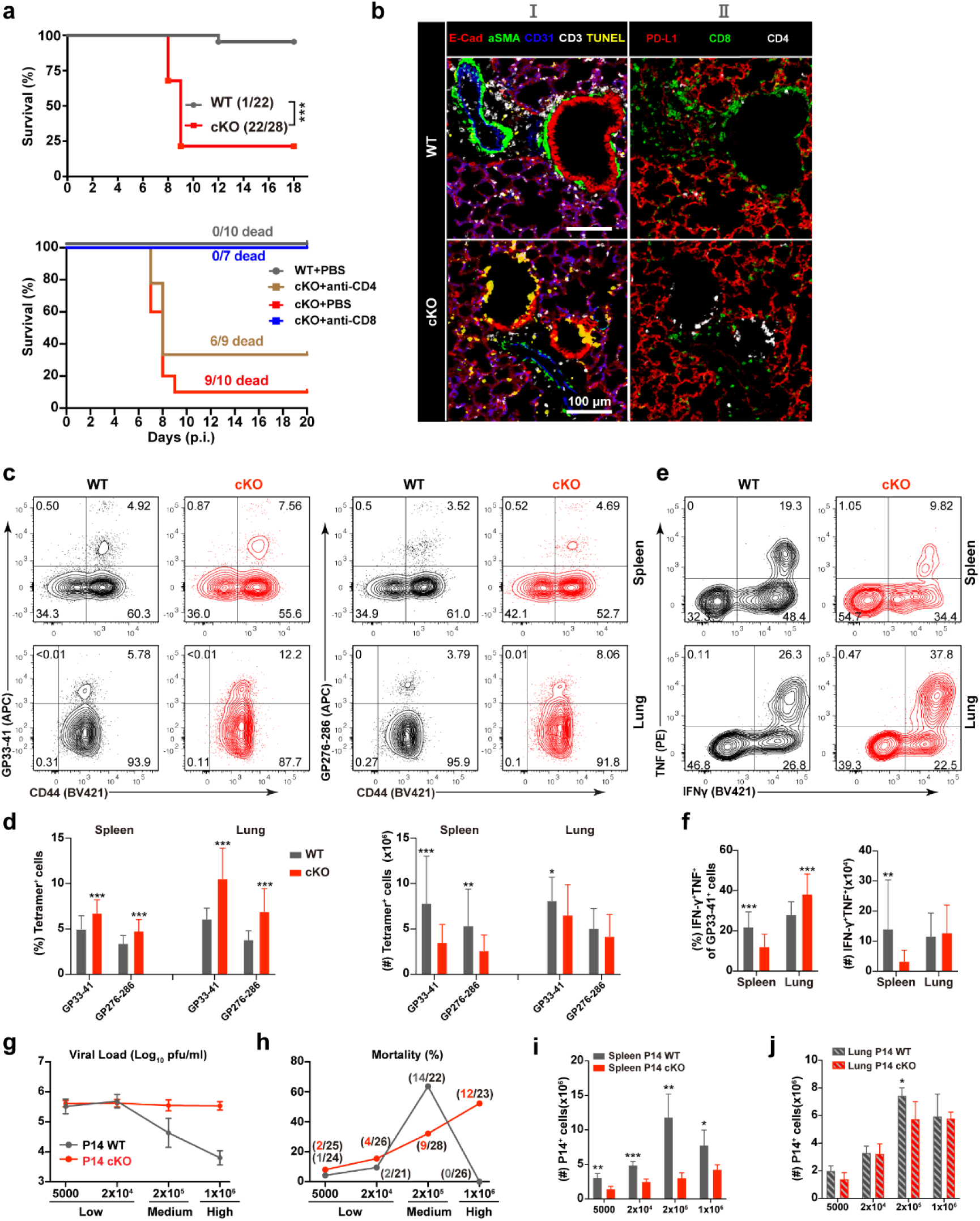
Themis T cell conditional knockout mice succumb to LCMV C13 infection. **a,** Survival of mice after LCMV C13 infection without (top) or with indicated cell depletion (bottom). **b,** Imaging Mass Cytometry of lung sections of mice at 8 dpi. Combinations of markers are shown as indicated. Bronchi were identified by adjacent staining of α-smooth muscle actin (αSMA) with E-cadherin (E-Cad), and blood vessels were identified by adjacent staining of αSMA with CD31, respectively. Scale bars, 100 μm. **c,d,** Quantitation of LCMV-specific CD8^+^ T cells. Mice were infected with LCMV C13 and analyzed at 8 dpi. Shown are representative FACS plots of tetramer staining, and summary of cell frequency and numbers as indicated. GP33-41 WT (n=33) and cKO (n=33); GP276-286 WT (n=28) and cKO (n=28). Data are pooled from multiple experiments. **e,f,** Cytokine measurement. CD8^+^ T cells were purified from infected mice and re-stimulated with GP33-41 peptide and GP33-41 tetramer^+^ cells were gated for analysis. WT (n = 27), cKO (n = 27). Data are pooled from multiple experiments. Shown are representative FACS plots and summary of cell frequency and number. **g-j,** Naive mice were transferred with indicated dose of P14 cells and then infected with LCMV C13 and analyzed at 8 dpi. (g) Virus titers determined by Q-PCR. (h) Mice mortality. Number of deceased mice out of total number of mice listed in brackets. Numbers of recovered P14 donor cells in the spleen (i) and lung (j). Data are pooled from three experiments (h). Experiments were performed 3 times (g, i, j), and representative results are shown. Error bars show SD. Statistical comparison of experimental groups was performed using unpaired Student’s t-test (d, f, i, j), or log-rank (Mantel-Cox) test for (a). NS, non-significant, * p < 0.05, ** p < 0.01, *** p <0.001.

To determine which type of T cells was responsible for the morbidity and mortality in cKO mice, we performed cell depletion and found that depletion of CD8^+^ T cells completely rescued mortality and lung injury in cKO mice, whereas depletion of CD4^+^ T cells only partially prevented mortality (Fig. 1a; Extended Data Fig. 1g), indicating that CD8^+^ T cells are the major mediators of casualties in cKO mice. When characterizing these CD8^+^ T cells, we found that, although the frequencies of LCMV-specific tetramer^+^ T cells were increased in the spleen and lung of cKO mice at 8 dpi, their numbers were mostly reduced compared with those of WT mice, especially in the spleen (Fig. 1c and d). Functionally, the proportion of GP33-41 tetramer^+^ cells co-expressing TNF and IFNγ in the spleen of cKO mice was lower than that in WT mice, but significantly higher than that in the lung, and the number of cells showed a similar trend (Fig. 1e and f). Elimination of GP33-41 peptide-loaded target cells was also impaired in cKO mice (Extended Data Fig. 1h), likely due to a reduction of GP33-41^+^ T cells in cKO mice. Together, these results showed that Themis-deficient LCMV-specific CD8^+^ T cells have unique properties, namely that they are defective in clonal expansion at the population level in the spleen (reduced numbers) but superior in effector function at the single-cell level in the lung (enhanced function). However, this led us to question whether these cells are sufficient to cause mortality in mice.

To answer the above question, we transferred different doses of P14 TCR transgenic cells generated in cKO or WT mice backgrounds (hereafter P14 cKO and P14 WT cells, see Methods) into C57BL/6 mice and then infected them with LCMV C13 (Extended Data Fig. 2a). Both P14 cell types recognize the GP33-41 peptide presented by H-2D^b^ and exhibited similar cytokine production (Extended Data Fig. 2b), cytotoxicity (Extended Data Fig. 2c), and proliferation (Extended Data Fig. 2d) in vitro, but P14 cKO cells showed a slight delay in activation^21^. Consistent with previous studies^24^, P14 WT cells reduced viral load in a dose-dependent manner, and mice receiving a medium dose of P14 WT cells showed significant mortality (Fig. 1g and h). In contrast, P14 cKO cells failed to control virus at all doses, and the mortality of recipient mice increased with increasing cell doses due to severe lung immunopathology (Fig. 1g, h, and Extended Data Fig. 2e). Interestingly, we found that the numbers of P14 cKO cells were also significantly reduced in the spleen but mostly comparable to those of P14 WT cells in the lung (Fig. 1i and j). Furthermore, we did not find mutant viruses in mice receiving high dose of P14 cKO cells, but we found previously reported GP35A mutant virus^25^ in mice receiving high dose of P14 WT cells (Extended Data Fig. 2f). These results suggested that Themis-deficient CD8^+^ T cells are inefficient to eliminate the virus but are sufficient to induce immunopathology.

### Themis deficiency promotes differentiation of effector T cells (T-eff) but inhibits exhausted T cell progenitors (T-pex)

To obtain clues about the abnormalities of P14 cKO cells, we performed scRNA-seq on co-transferred P14 cells to avoid interference from environmental factors (Extended Data Fig. 3a). We analyzed cells at 5 dpi because the effects observed at 8 dpi ought to be traced back to earlier transcriptional changes. We combined P14 WT and P14 cKO samples from the spleen and lung for t-SNE analysis and identified 10 cell clusters by unsupervised clustering (Fig. 2a) that exhibited extreme tissue and genotype specificity (Extended Data Fig. 3b). We first evaluated differential gene expression between the two P14 cell types and found that RNA expression of *Pdcd1* (encoding PD-1) and *Tox* was significantly downregulated in P14 cKO cells, whereas expression of the cytotoxic molecules *Gzmb*, *Gzmk*, and *Gzma* was upregulated, regardless of tissue type (Extended Data Fig. 3c). For cluster characterization, we found that clusters 6 and 7 had similar gene expression profiles (Extended Data Fig. 4) and express *Sell* (encoding CD62L), and *Tcf7* (encoding TCF-1) (Fig. 2b), indicating that they are exhausted T cell progenitors (T-pex). Notably, clusters 6 and 7 were mainly present in the spleen and lung, respectively, and both consisted of half P14 cKO cells and half P14 WT cells (Fig. 2c and d). Clusters 0 and 1 also exhibited similar gene expression profiles (Extended Data Fig. 4) and expressed high level of *Klrg1* and *Cd69* (Fig. 2b). They were dominant in P14 cKO and P14 WT cells in the lung (Fig. 2c and d), indicating that they are lung-localized effector T cells (T-eff). Notably, cluster 1 expressed higher levels of *Tox* than cluster 0 (Fig. 2b). Clusters 2, 3, and 5 all expressed high levels of *Gzmb* and the proliferation marker *Mki67* (Fig. 2b), but the former was dominant alone in P14 WT cells in the spleen, whereas the latter two were dominant together in P14 cKO cells in the spleen (Fig. 2c and d), indicating that they are developing T-eff cells.

**Fig. 2.**
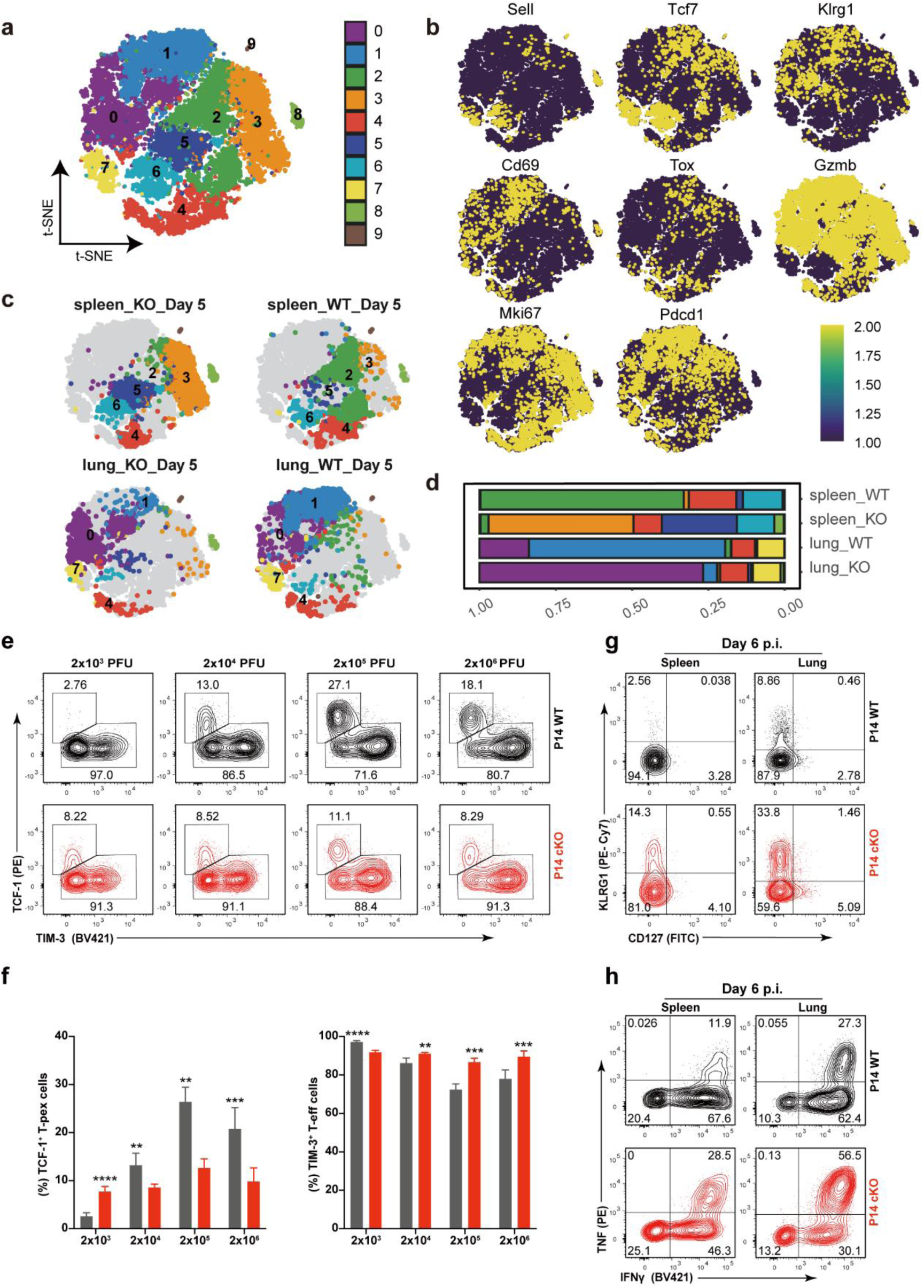
Themis deficiency promotes T-eff cell differentiation but inhibits T-pex cell differentiation. **a-d,** scRNA-seq analysis. The following cell counts were used: Lung/cKO = 4684 cells, Lung/WT = 6460 cells, Spleen/cKO = 8289 cells, Spleen/WT = 5779 cells. (a) t-SNE projections of pooled cells from all four samples. Analysis by Seurat v.4 revealing 10 distinct clusters (labeled 0–9), each distinguished by a unique color code. (b) Single-cell transcript levels of key genes, visualized in t-SNE plots, with expression color-coded: black (non-expressed), yellow (expressed). The distribution (c) and proportion (d) of cell clusters in the spleen and lung are shown as indicated. **e,f,** Expression of TCF-1 and TIM3 in co-transferred P14 cells after recovery from recipient mice at 5 dpi with indicated virus doses. Shown are representative FACS plots and summary of the frequency of TCF-1^+^ T-pex and TIM3^+^ T-eff cells. **g,h,** Expression of KLRG1 and CD127 (g) and IFNγ and TNF (h) in co-transferred P14 cells at 6 dpi. Shown are representative FACS plots. Experiments were performed 2 times (e, g, h). Error bars show SD. Statistical comparison of experimental groups was performed using paired Student’s t-test (f). * p < 0.05, ** p < 0.01, *** p <0.001, **** p <0.0001.

Our scRNA-seq data are consistent with current models that suggest that CD8^+^ T cells initially bifurcate into T-eff and T-pex during the early stages of chronic LCMV infection^6,26–29^. To verify the above findings at the cellular level, we further analyzed two types of P14 cells. Recent studies have shown that T-pex and T-eff can be distinguished phenotypically as early as 5 dpi in chronic LCMV infection, and that their differentiation depends on the amounts of viral antigens encountered^6^. As previously reported, we found that P14 WT cells branched into T-pex (TCF-1^+^TIM3^−^) and T-eff (TCF-1^−^TIM3^+^), and the proportion of T-pex increased roughly as the virus dose increased (Fig. 2e and f). However, this trend was not obvious in P14 cKO cells. In addition, in P14 cKO cells, the proportion of T-pex was mostly reduced, while the proportion of T-eff was increased (Fig. 2e and f). We also used KLRG1, a typical T-eff marker^26,30,31^, to confirm that a large number of T-eff appeared early in P14 cKO cells, while such cells were rare in P14 WT cells (Fig. 2g). Correspondingly, more P14 cKO cells co-expressed TNF and IFNγ than P14 WT cells (Fig. 2h). Together, these results suggest that Themis deficiency promotes T-eff differentiation but inhibits T-pex differentiation.

### Ordered regulation of TCR and PD-1 signaling by Themis in chronic LCMV infection

Although our above results in P14 WT cells supported the idea that high viral loads and the resulting strong TCR signaling favor T-pex differentiation over T-eff^6^, we do not know the situation in P14 cKO cells. We hypothesized that disruption of Themis would alter early TCR signaling, thereby affecting the process of T cell exhaustion. Furthermore, Themis deficiency resulted in increased cytokine production in CD8^+^ T cells, a sign of disrupted T cell exhaustion. To accurately investigate this, we utilized Nur77^GFP^ transgenic mice^32^ and generated P14 WT/Nur77^GFP^ and P14 cKO/Nur77^GFP^ cells (see Methods), whose TCR signaling can be reported by GFP expression. To examine early TCR signaling, we established a cell coculture system in which P14 WT/Nur77^GFP^ and P14 cKO/Nur77^GFP^ cells carrying different congenic markers were mixed with splenocytes from C13-infected mice to best mimic the in vivo situation. We found that the proportion of GFP^+^ P14 cKO/Nur77^GFP^ cells was lower than that in P14 WT/Nur77^GFP^ cells from 6 to 24 hours of culture (Fig. 3a), indicating that Themis deficiency impaired early TCR signaling. Next, we performed in vivo analysis by adjusting the number of transferred cells accordingly (Extended Data Fig. 5a). Remarkably, the ratio of P14 cKO/Nur77^GFP^ cells to P14 WT/Nur77^GFP^ cells changed drastically over time, with P14 cKO/Nur77^GFP^ cells predominating early in infection (3 and 4 dpi) but becoming subdominant (5 dpi) and almost disappearing at later stages (Fig. 3b, and Extended Data Fig. 5b). These results suggest that the initial generation of P14 cKO/Nur77^GFP^ effector cells in the spleen and their migration to the lung is much more vigorous than those of P14 WT/Nur77^GFP^ cells, but this trend rapidly declines with infection.

**Fig. 3.**
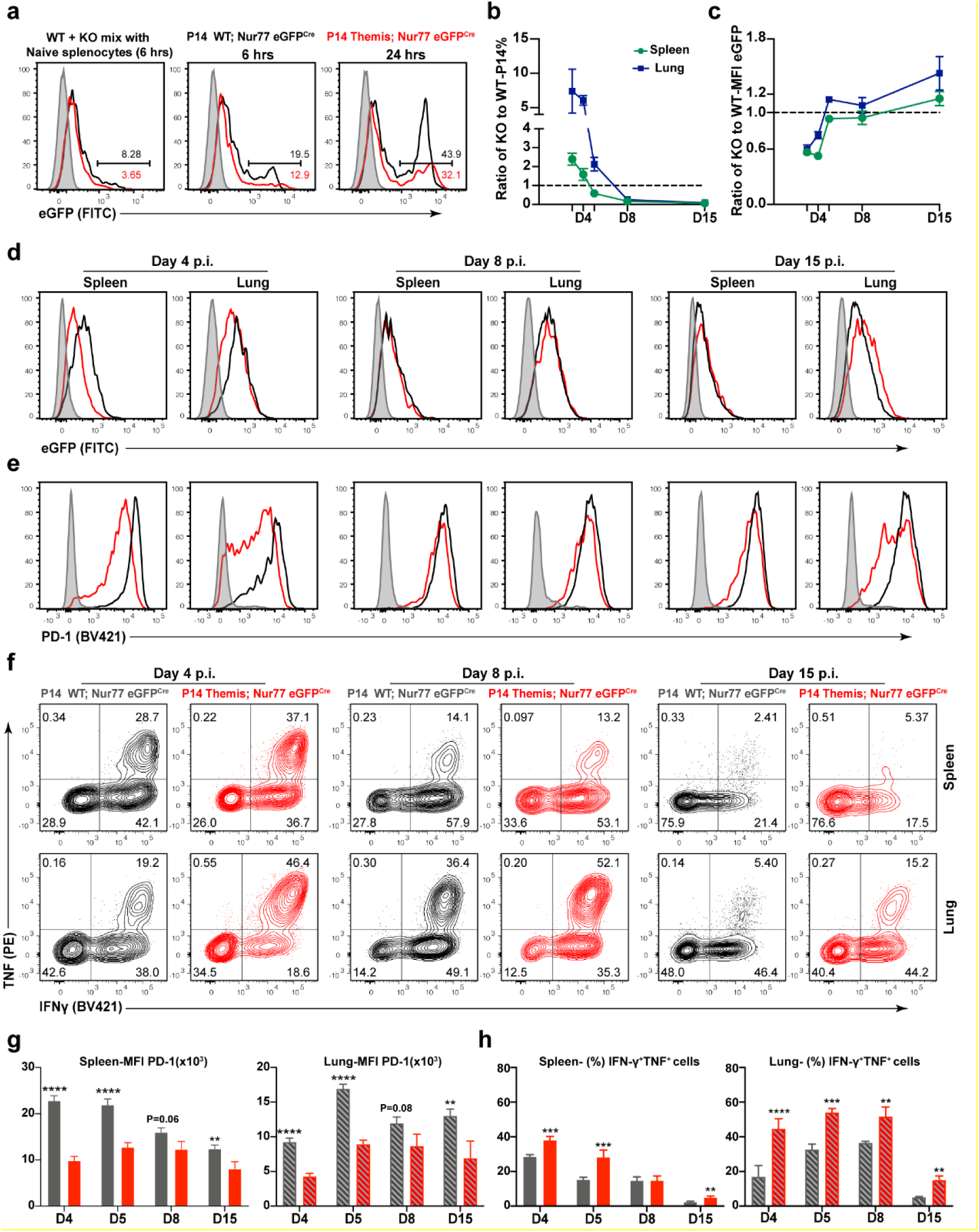
Orderly regulation of TCR and PD-1 signaling by Themis. **a,** Naive P14 WT/Nur77^GFP^ and P14 cKO/Nur77^GFP^ cells with different congenic markers were purified and co-cultured with splenocytes derived from LCMV C13-infected mice. GFP expression level are measured at 6 and 24 hours. **b,** The ratio of P14 cKO/Nur77^GFP^ cells to P14 WT/Nur77^GFP^ cells at indicated time points in the spleen and lung. **c,** The ratio of GFP level of P14 cKO/Nur77^GFP^ cells to P14 WT/Nur77^GFP^ cells at indicated time points in the spleen and lung. **d-f,** Representative FACS plots of GFP (d), PD-1 (e), and IFNγ and TNF (f) expression in P14 cells as indicated. **g,h,** Summary of PD-1 expression (g) and IFNγ and TNF co-expression (h) in P14 cells as indicated. All experiments were performed 2 times. Statistical comparison of experimental groups was performed using paired Student’s t-test (g, h) Error bars show SD. * p < 0.05, ** p < 0.01, *** p <0.001, **** p <0.0001.

We next examined TCR signaling in P14 cells by GFP levels and found that the ratio of GFP levels in P14 cKO/Nur77^GFP^ cells to those in P14 WT/Nur77^GFP^ cells increased over time, especially in the lung (Fig. 3c). Specifically, the GFP level in P14 cKO/Nur77^GFP^ cells was lower than that in P14 WT/Nur77^GFP^ cells at 3 and 4 dpi, became comparable at 5 and 8 dpi, and exceeded the latter at 15 dpi (Fig. 3d, and Extended Data Fig. 5c). Consistent with reports that PD-1 expression is induced by TCR stimulation^33^, PD-1 expression in P14 cKO/Nur77^GFP^ cells varied with the intensity of TCR signals. Specifically, PD-1 expression in P14 cKO/Nur77^GFP^ cells was much lower than that in P14 WT/Nur77^GFP^ cells at 3 and 4 dpi, but their difference gradually narrowed from 5 to 8 dpi (Fig. 3e, and Extended Data Fig. 5d), corresponding to the relative elevation of TCR signal in these cells. Interestingly, PD-1 expression in P14 cKO/Nur77^GFP^ cells was significantly reduced again at 15 dpi (Fig. 3e), suggesting that Themis may also have long-term effects in T cell exhaustion. For cytokines, P14 cKO/Nur77^GFP^ cells co-expressed IFNγ and TNF more than P14 WT/Nur77^GFP^ cells in all conditions except in the spleen at 8 dpi (Fig. 3f, and Extended Data Fig. 5e). When the above changes were plotted and compared, we found a clear relationship: high PD-1 expression in P14 WT/Nur77^GFP^ cells resulted in low cytokine production, whereas low PD-1 expression in P14 cKO/Nur77^GFP^ cells resulted in high cytokine production in these cells (Fig. 3g and h). These results collectively suggest that Themis promotes TCR signaling to induce PD-1 expression, which in turn inhibits cytokine production. In the absence of Themis, TCR signaling is attenuated, as is PD-1 expression and its inhibitory function. Importantly, these results demonstrate that early TCR signaling has a significant and lasting impact on the development of T cell exhaustion.

### Themis is required for maintenance of terminally differentiated exhausted T cells (T-ex)

Our observation that PD-1 expression in P14 cKO cells was reduced abruptly again late in infection (Fig. 3e) suggests that Themis may have long-term effects on T cell exhaustion. To further explore this, we extended our cell co-transfer experiments to 30 dpi, when T cell exhaustion is fully established (Extended Data Fig. 6a). Studies have shown that after initial differentiation, T-pex cells continue to self-renew and give rise to terminally differentiated exhausted T cells (T-ex) as infection persists, which can be distinguished by the expression of transcription factors TCF-1 and TOX^26,28,30,34,35^. We found that T-pex cells (TCF-1^+^TOX^+^) were consistently present in P14 WT cells in the spleen throughout infection but were virtually absent in P14 cKO cells and could only be transiently detected in both cell types in the lung (Fig. 4a and b). As expected, T-ex cells (TCF-1^-^TOX^+^) were also severely reduced in P14 cKO cells, especially at late stage (Fig. 4a and b). Because TOX has been shown to promote the expression of PD-1 and other co-inhibitory molecules^30,36,37^, we examined them and found that only PD-1 expression was significantly reduced over time in P14 cKO cells (Extended Data Fig. 6b and c), whereas other co-inhibitory molecules did not decrease or only slightly decreased (Extended Data Fig. 6d and e).

**Fig. 4.**
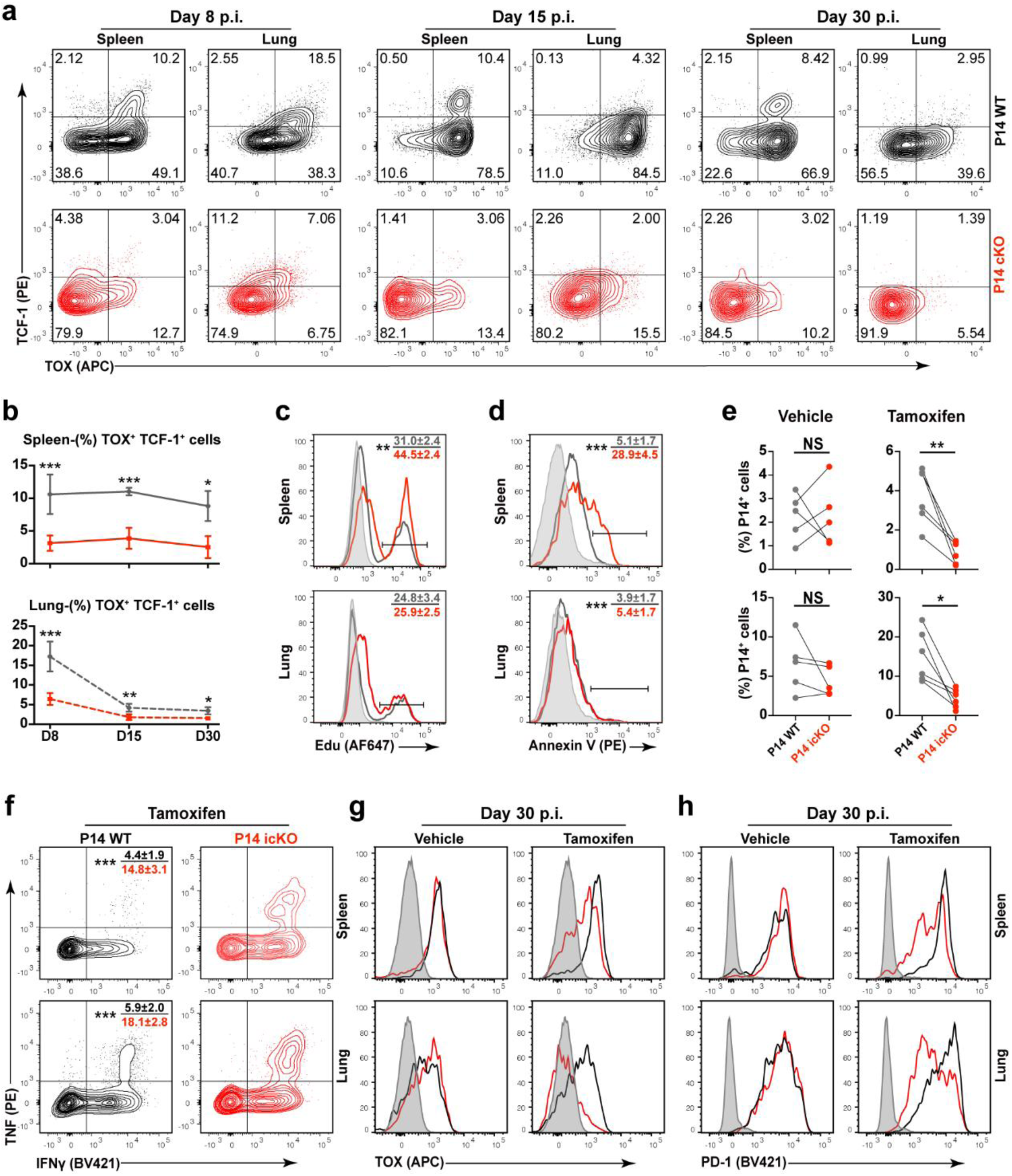
Themis is required for the maintenance of T-ex cells. **a,b,** Expression of TCF-1 and TOX in co-transferred P14 cells at indicated time points. Shown are representative FACS plots (a) and summary of the frequency of TCF-1^+^TOX^+^ cells (b). **c,** In vivo cell proliferation. Mice receiving P14 cells were given EdU and analyzed at 8 dpi. Shown are representative FACS plot with mean ± SD listed in the plot. **d,** Cell death analysis. P14 cells were analyzed ex vivo by Annexin V staining at 8 dpi. Shown are representative FACS plot with mean ± SD listed in the plot. **e-h,** Analysis of P14 cells at 30 dpi after two rounds of tamoxifen treatment. (e) The proportion of P14 cells. (f) Expression of IFNγ and TNF. Shown are representative FACS plots. Expression of TOX (g) and PD-1 (h). Shown are representative histogram overlays of P14 WT and P14 icKO cells. Gray shadows represent splenocytes from naive mouse, used as control. Experiments were performed two times. Statistical comparison of experimental groups was performed using paired Student’s t-test (b, e) Error bars show SD. (NS, non-significant, * p < 0.05, ** p < 0.01, *** p <0.001, **** p <0.0001).

One recurring unanswered question in our transfer experiments is the rapid loss of P14 cKO cells, as occurred here (Extended Data Fig. 7a) and previously (Figs. 3b, and 1I). To understand this, we examined the residual P14 cells and found that a larger proportion of T-eff cells (Extended Data Fig. 7b and c) as well as IFNγ and TNF co-expressing cells (Extended Data Fig. 7d and e) were consistently present in P14 cKO cells. Since T-effs are known to be both proliferative and transient, we found that P14 cKO cells in the spleen proliferated 1.4-fold faster and died 5.6-fold more rapidly than P14 WT cell, whereas changes in the lung were minimal (Fig. 4c, d). Together, these results reinforce and extend our previous findings (Fig. 2e), showing that Themis deficiency impairs T-pex differentiation in a long-term irreversible manner. Furthermore, the rapid turnover of Themis-deficient T-effs in the spleen was primarily responsible for the clonal expansion defects seen in infected cKO mice and in P14 cKO transfer models.

Although we have observed that Themis deficiency leads to severe defects in T-ex cells (Fig. 4a), this is most likely a secondary effect due to the primary defect in T-pex differentiation. To test whether Themis directly regulates the long-term maintenance of T-ex cells, we used a tamoxifen-mediated gene deletion strategy to rapidly delete *Themis* in P14 Themis^flox/flox^ ERT2-Cre cells^20^ (hereafter P14 icKO, see Methods), bypassing the early infection stage and keeping T-pex intact (Extended Data Fig. 8a). Baseline analyses showed that P14 icKO and P14 WT cells exhibited comparable cell number and cytokine production (Extended Data Fig. 8b and c) as well as TOX and PD-1 expression (Extended Data Fig. 8d and e). We confirmed that Themis was effectively deleted at 18 dpi (Extended Data Fig. 8f). Although there were no quantitative differences between P14 WT and P14 icKO cells (Extended Data Fig. 8g), the proportion of P14 icKO cells co-expressing TNF and IFNγ was increased (Extended Data Fig. 8h), and TOX and PD-1 expression was slightly reduced in P14 icKO cells (Extended Data Fig. 8i and j). At 30 dpi, the proportion of P14 icKO cells decreased dramatically compared with P14 WT cells, whereas there was no difference between the two cell types treated with vehicle (Fig. 4e). Furthermore, the proportion of P14 icKO cells co-expressing TNF and IFNγ was significantly higher than that of P14 WT cells (Fig. 4f), while the expression of TOX and PD-1 was significantly reduced in P14 icKO cells (Fig. 4g and h). Taken together, these results demonstrated that Themis directly promotes the long-term maintenance of T-ex cells and their function.

### Themis is an essential mediator of PD-1 signaling

Although decreased PD-1 expression could explain the increased cytokine production in P14 cKO cells in most cases, this may be insufficient for the situation in the lung at 8 dpi (Fig. 3g and h). More importantly, in cKO mice infected with LCMV C13, endogenous Themis-deficient virus-specific CD8^+^ cells expressed variable levels of PD-1 compared to WT cells (Fig. 5a), but consistently produced more cytokines in the lung (Fig. 1e and f), suggesting that Themis deficiency may disrupt PD-1 signaling independent of impairing PD-1 expression.

**Fig. 5.**
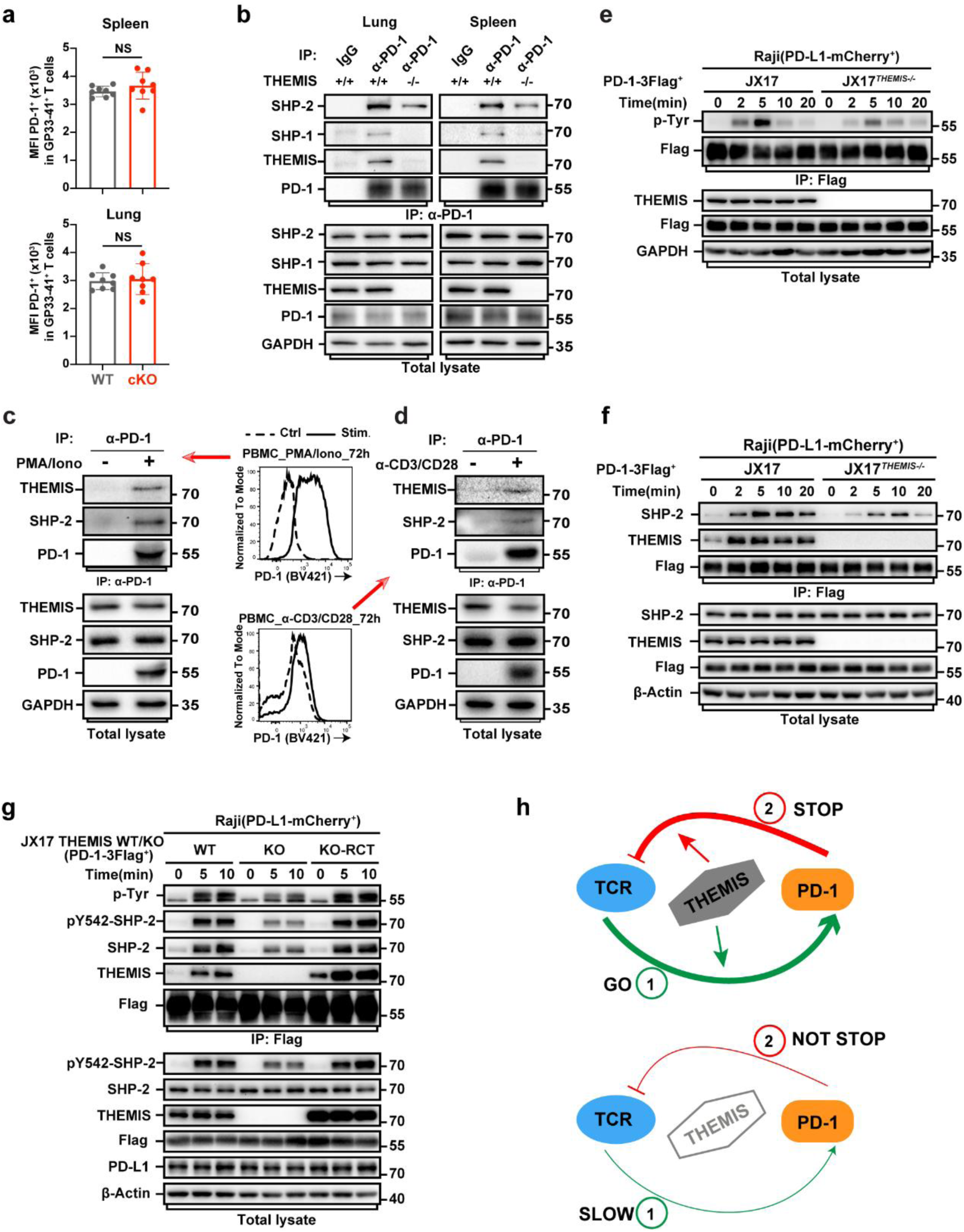
Themis is required for PD-1 signaling. **a,** PD-1 expression in tetramer GP33-41^+^ T cells in WT and cKO mice at 8 dpi. MFI, mean fluorescence intensity. **b,** Themis and PD-1 interaction. Primary CD8^+^ T cells were FACS sorted from the lung and spleen of WT and cKO mice at 8 dpi. PD-1 was immunoprecipitated from cell lysates and detected for interactions with indicated proteins. **c,d,** T cells were purified from healthy donors’ peripheral blood mononuclear cells (PBMCs), and stimulated with PMA and Ionomycin (c) or anti-CD3/CD28 (d) for 3 days and viable cells were FACS sorted. The induced expression of PD-1 were verified by FACS. The interaction between endogenous PD-1, THEMIS, and SHP2 were determined by co-immunoprecipitation as indicated. Shown are one of two experiments. **e,** THEMIS deficiency diminished PD-1 phosphorylation. Jurkat-PD-1-3Flag cells were stimulated with Raji-PD-L1 cells. Cell lysates were immunoprecipitated with anti-Flag Sepharose beads and immunoblotted with anti-phosphotyrosine (p-Tyr) antibody. **f,** THEMIS deficiency impaired PD-1 and SHP2 binding. The same experimental procedure as described in (e), except with antibodies to SHP2 and THEMIS. **g,** THEMIS was both necessary and sufficient for PD-1 signaling. JX17 THEMIS KO-RCT cells were generated by reconstituting JX17 THEMIS KO cell line with recombinant THEMIS. Cells were stimulated as in (e). Cell lysates were immunoprecipitated with anti-Flag Sepharose beads and immunoblotted with indicated antibodies. **h,** Themis working model as described in the text. Experiments depicted in (a) was performed 3 times with five to eight mice per group. Experiments were performed two or three times (b-g) and representative results are shown. Statistical comparison of experimental groups was performed using unpaired Student’s t-test (a) Error bars show SD. (NS, non-significant, * p < 0.05, ** p < 0.01, *** p <0.001, **** p <0.0001).

To test the above hypothesis, we first sought to verify whether Themis binds to PD-1 in primary CD8^+^ T cells extracted from LCMV-infected mice. We found that Themis, SHP2, and to a much lesser degree SHP1, were co-precipitated with PD-1 (Fig. 5b). Similar results were obtained in human PBMCs after pretreatment to induce PD-1 expression (Fig. 5c and d). Notably, the interaction between PD-1 and SHP2 was significantly reduced in CD8^+^ T cells obtained from cKO mice (Fig. 5b), indicating that Themis is important for PD-1-SHP2 association. To further investigate how THEMIS mediated PD-1 signaling, we generated a THEMIS-deficient Jurkat cell line (JX17^THEMIS-/-^)^38^, and used it in our previously developed Raji-Jurkat stimulation system^39^. We found that in the absence of THEMIS, phosphorylation of PD-1 was severely impaired (Fig. 5e), which is responsible for the reduced recruitment of SHP2 by PD-1 (Fig. 5f), as previously reported^40^. Finally, we engineered a rescue cell line (KO-RCT) in which THEMIS expression was restored in the above JX17^THEMIS-/-^ cells. In this scenario, we found that the phosphorylation of PD-1 was restored, as was the binding of PD-1 to SHP2. Moreover, while the phosphorylation of SHP2 was significantly reduced in the absence of THEMIS, it was recovered upon THEMIS reconstitution (Fig. 5g). These results suggest that in addition to playing a role in promoting TCR signaling and PD-1 expression, Themis is also critical for PD-1 phosphorylation and signaling. We therefore propose a model whereby Themis sequentially promotes TCR and PD-1 signaling, one for T cell activation and one for T cell inhibition, forming a “GO-STOP” normal regulatory pattern. In the absence of Themis, not only is TCR signaling impaired, but PD-1 signaling is also disrupted, resulting in a “SLOW but NOT STOP” failure pattern (Fig. 5h).

## Discussion

Counterintuitively, CD8^+^ T cells lacking Themis are defective in both homeostasis (Extended Data Fig. 9a, Box I)^19^ and clonal expansion, but still cause lung damage and mortality in mice during chronic LCMV infection. We speculate that without the rapid turnover of these cells, the “enhanced function” property of Themis-deficient CD8^+^ T cells (Extended Data Fig. 9a, Box II and III) may not be sufficient to compensate for the “reduced numbers” effect, because rapid turnover actually promotes the accumulation of large numbers of virus-specific CD8^+^ T cells. The “enhanced function” of Themis-deficient CD8^+^ T cells can be explained by the “slow but not stop” model mentioned above (Fig. 5h). The dangerousness of this model is maliciously amplified when the virus persists (Extended Data Fig. 9b) but is harmless in acute LCMV infection when the virus is effectively cleared^21^ (Extended Data Fig. 9c).

Although Themis not only promotes PD-1 expression but also mediates it’s signaling, Themis deficiency is not an equivalent to PD-1 deficiency. We found that immunopathology in cKO mice was restricted to the lung, in contrast to the multi-organ damage reported in PD-1 KO mice^41^. Furthermore, while one million P14 cKO cells were unable to control virus in LCMV-infected recipient mice, 100-fold smaller number of P14 PD-1 KO cells was able to do so^41^. Similarly, while large numbers of P14 cKO cells are required to cause severe mortality in LCMV C13-infected mice, only a few hundred P14 PD-1 KO cells are needed^42^. These differences are most likely because PD-1 deficiency results in both numerical and functional gain, whereas Themis deficiency results in functional gain only (Extended Data Fig. 9b). The exact molecular mechanisms by which Themis regulates PD-1 expression and signaling remain unclear and warrant further investigation^43^.

Previous studies have shown that the degree of TCR engagement determines the differentiation of CD8^+^ T cells in the early stage of chronic LCMV infection^6^. We confirmed this finding and further demonstrated that intrinsic TCR signaling perturbations, such as Themis disruption, have profound effects on every aspect of T cell exhaustion (Extended Data Fig. 9a). This finding highlights the possibility that in addition to manipulating immune checkpoints, Themis and other signaling molecules could also be targeted to rewire TCR signaling and reverse or enhance T cell exhaustion, as has been attempted in CAR-T cells^44^.

## Methods

### Mice

Themis^flox/flox^-dLck-cre mice were generated at the National University of Singapore and re-derived at Xiamen University before a colony was established. All mouse experiments were approved by the Institutional Animal Care and Use Committee of Xiamen University. For all experiments, mice of 8 to 12 weeks old were used. P14 Themis^flox/flox^-dLck-Cre Rag1^-/-^ mice were generated by breeding Themis^flox/flox^-dLck-Cre mice with P14 TCR transgenic mice and Rag1^-/-^ mice (Jackson Laboratory Strain # 002216). P14 Themis^flox/flox^-ERT2-Cre mice were generated by breeding Themis^flox/flox^ mice with Rosa26-CRE^ERT2^ mice (Jackson Laboratory Strain #008463). P14 Themis^flox/flox^Nur77^GFP^-Cre mice were generated by breeding P14 Themis^flox/flox^ mice with Nur77^GFP^-Cre mice (Jackson Laboratory Strain #018974). In P14 Themis^flox/flox^-Nur77^GFP^-Cre mice, *Themis* is deleted in the thymus, which reduces the number of developing P14 cells. We confirmed that the residual P14 cells in P14 Themis^flox/flox^-Nur77^GFP^-Cre mice were still functional in vivo in transfer experiments. In addition, for each transfer experiment, we purified naive P14 cells from P14 Themis^flox/flox^-Nur77^GFP^-Cre mice or control P14-Nur77^GFP^-Cre mice and ensured that both cell types express the same or similar levels of GFP.

### Plasmids and reagents

Human PD-1 was sub-cloned into a modified pLV lentiviral vector with a C-terminal 3×Flag epitope tag for stable expression in Jurkat T cells. All plasmids were verified by DNA sequencing. SEE super-antigen (#ET404) was purchased from Toxin Technology.

### Antibodies

For western blots, anti-SHP1(#3759), anti-phospho-SHP1(Y564) (#8849S), anti-SHP2 (#3397), anti-phospho-SHP2(Y542) (#3751S), anti-p44/42 MAPK(Erk1/2) (#9107S), anti-phospho-p44/42 MAPK (T202/Tyr204) (#4370S), anti-human PD-1 (clone D4W2J) and anti-mouse PD-1 (D7D5W) were purchased from Cell Signaling Technology; Anti-Themis (#ab126771) was purchased from Abcam; Monoclonal anti-phosphotyrosine clone PY-20 (#P4110) was purchased from Sigma. Anti-PD-L1 (#66248-1-Ig), anti-beta Actin (#66009-1-Ig), anti-GAPDH (60004-1-Ig), and anti-Flag (#20543-1-AP) were purchased from Proteintech. For mouse PD-1 immunoprecipitation, rat anti-mouse PD-1 monoclonal antibody was provided by Dr. Chenghao Huang. This antibody was tested and validated before use. For Flag immunoprecipitation, anti-Flag M2 affinity Gel (#A2220) were obtained from Sigma-Aldrich.

### Antibody Conjugation

Lanthanide metal-conjugated antibodies used in Imaging mass cytometry were listed in Extended Data Fig. 1c. Un-conjugated antibodies were purchased and labeled with selected isotopes using the MaxPar X8 labeling kit from Fludigm according to the manufacturer’s instruction. Briefly, the un-conjugated antibody was partially reduced with TCEP buffer and then incubated with the metal-loaded MaxPar X8 polymer. The metal-labeled antibody was recovered by an exclusion filter and stored in the antibody stabilizer before long-term storage at 4°C. To optimize the detection of positive populations and minimize the background, serial dilution staining experiments were performed.

### Generation of knockout and overexpression cell lines

A Jurkat T cell line that stably expressing Cas9 protein, JX17, was kindly provided by Dr. Haopeng Wang (Shanghai University of Science and Technology). Jurkat and JX17 cells were cultured in complete RPMI 1640 medium supplemented with 10% FBS and antibiotics at 37 ℃ in a 5% CO_2_ incubator. Knockout cell line was generated by CRISPR/Cas9 system. gRNA targeted *THEMIS* gene candidate sequences are ATAGCAATGGCATTATCAC, and TACCCAGGGTTCTAGAAATC. The knock-out construct is based on pLKO-EGFP vector. After cloning, JX17 cells were electroporated with gRNA-expressing plasmid. Two days later, GFP-positive cells were sorted by flow cytometry on the BD Aria III. Single cell clones of sorted cells were obtained by limiting dilution. Knockout clones were identified by sequencing of the PCR fragments, and confirmed by western blot using THEMIS antibody (Abcam; ab126771). For PD-1 overexpression, HEK293T cells were transfected with packaging plasmids and pLV-based vector by calcium phosphate transfection. Virus-containing supernatants were collected 48 hrs later and used to infect Jurkat T cells.

### Jurkat T cells stimulation assay

Raji B cells were pre-incubated with 30 ng/mL SEE super-antigen for 30 min at 37℃. Jurkat T cells were rested in serum free RPMI medium at 37 °C for 3-5 hrs. Then, Jurkat T cells expressing hPD-1-3×Flag were stimulated with Raji B cells expressing PD-L1. The cells were lysed with lysis buffer (20 mM Tris-Hcl, 150 mM NaCl, and 1 mM EDTA) containing 1 % NP-40 with protease and phosphatase inhibitor cocktail, and immunoprecipitated with anti-Flag antibody.

### Immunoprecipitation and immunoblotting

For Jurkat T cells, immunoprecipitations were performed using anti-Flag M2 beads, or anti-PD-1 antibody with Protein A/G agarose beads. Western blotting of cell lysates and immunoprecipitates were performed using anti-Flag, anti-SHP2, anti-phospho-SHP2, anti-THEMIS, anti-p-Tyr, and anti-PD-L1. For primary cells, mouse CD8^+^ T cells were prepared from lungs and spleens of infected mice. Briefly, lungs were harvested from mice, mechanically sheared into small pieces, and digested in RPMI containing collagenase (type 4) (Aladdin; 1 mg/mL), and incubated for 1 hr at 37 °C. After digestion, cells were passed through a 100-μm filter, and red blood cells were lysed by ACK lysis buffer. Lung and spleen CD8^+^ T cells were further purified by FACS sorting. Purified primary CD8^+^ T cells were finally re-suspended with lysis buffer. For immunoprecipitation, cell lysate was centrifuged at 13,000 rpm for 10 min at 4 °C to generate cleared supernatant, and protein A/G beads were added into the supernatant followed by rotation at 4 °C for 45 min. Anti-PD-1 was added into supernatant and incubated at 4 °C overnight with rotation. Then protein A/G beads were added and rotated for 2 hrs at 4 °C. The beads were washed with immunoprecipitation buffer three times at 4 °C and then mixed with an equal volume of 2×SDS sample buffer for western blot. When required, lung-infiltrated inflammatory cells were purified from prepared cell suspension by centrifugation in PBS buffered Percoll (GE Healthcare Life Sciences; # 17089101).

### LCMV infection and virus titer test

Stocks of lymphocytic choriomeningitis virus (LCMV) strains Clone 13 were prepared by serial passage in BHK-21 cells. For infection, 2×10^6^ PFU was injected intravenously per mouse. For virus titer measurement in the blood, mouse serum was collected, and 15 μL of serum was serially diluted by 10-fold for plaque assays on Vero cells. For virus titer measurement in the organs, lungs and livers were collected after dissecting mice, and tissues were frozen at -80 °C. At the time of assay, tissues were thawed, weighed, and subsequently homogenized in medium. Supernatants were prepared by centrifugation, and 15 μL of the supernatant was serially diluted by 10-fold for plaque assays.

### Cell depletion

When cell depletion is needed, mice were treated intraperitoneally with anti-CD4 monoclonal antibody YTS191.1 (BioXCell; BE0119-25MG) on days -1, 3, 5, and 7 before and post LCMV infection (200 μg per treatment). Similar doses and schedules were used for CD8^+^ T cells with the monoclonal antibody YTS169 (BioXCell; BE0117-25MG).

### Measurement of cytokine levels in BAL

BAL cytokine concentration was measured using MILLPLEX MAP Mouse Cytokine/Chemokine Magnetic Bead Panel-Premixed 25 Plex-Space Saver (Bulk) Packaging kit (Merck; MCYTMAG70PMX25BK), and the Luminex® analyzers Luminex® 200TM.

### Vascular Leakage Assay

At the time of assay, mice were intravenously injected with 200 μL of Evan’s Blue dye (10 mg/ml in PBS). Twenty minutes post dye injection, mice were sacrificed and perfused by intracardial injection of 15 mL of PBS, and lungs were collected.

### Histology

Lungs were harvested from infected mice, placed in PBS-buffered formalin, and blocked in paraffin. 5-μm tissue sections were stained with hematoxylin and eosin. Microscopic images were captured on Motic VM1 Microscopic digital slice scanning system.

### Immunofluorescence

Isolated PBS-perfused lungs were fixed with 4% Paraformaldehyde (Sigma-Aldrich) at 4°C overnight, under constant agitation. Then dehydrated in 30% sucrose before embedded with Tissue-Tek OCT compound (Sakura). Sections of 7 μm were permeabilized and using Cell Apoptosis Detection Kit (Boster Biological Engineering Co., Ltd., MK1019) for TUNEL staining according to the manufacturer’s instructions. After staining, sections were blocked in PBS containing 3 % BSA and 0.4 % TritonX-100 (Sigma-Aldrich) for 1 hr at room temperature followed by incubating with antibodies diluted in blocking buffer overnight at 4°C. Following antibodies were used: Anti-wide spectrum Cytokeratin antibody (Abcam; ab9377) and rat anti-mouse CD31 antibody (Abcam; ab7388). Sections were washed with PBS and nuclei were stained with 10 μg/mL DAPI (Invitrogen; D3571), and co-incubating with second antibodies containing Goat anti-Rat IgG (H+L) Alexa Fluor 594 (Invitrogen; A11007) and Donkey anti-Rabbit IgG (H+L) Alexa Fluor 647(Invitrogen; A21447) diluted in blocking buffer for 1 hr at room temperature and then washed with PBS followed by mounted with ProLong Gold Antifade Mountant (Molecular probes; P36934). All images were acquired using a Leica SP8 Laser Scanning Confocal Microscopy and analyzed by LAS X software.

### Tetramer, cell surface and intracellular staining, and cell sorting

For tetramer staining, cells were incubated with MHC-I tetramers H-2D^b^/GP33–41 or H-2D^b^/GP276–286 before any additional surface staining. All tetramers were made by the reagent lab of Cancer Research Center of Xiamen University. For cell surface staining, cells were incubated with antibody cocktail at 4 °C for 30 min. Antibody cocktail was prepared by diluting antibodies in FACS buffer (PBS supplemented with 0.5 % BSA and 0.01 % azide). In the cocktail, following antibodies were used: KLRG1(2F1), TIM-3 (RMT3-23), LAG-3 (C9B7W), IFNγ (XMG1.2), TNF (MP6-XT22), CD45.1 (A20), CD45.2 (104) from eBioscience; CD8α (53-6.7), CD44 (IM7), CD244.2 (2B4), CD160 (7H1), TIGIT (1G9), CD127 (A7R34), PD-1 (29F.1A12); CD90.1 (OX-7), CD90.2 (53-2.1) from Biolegend; TOX (REA473) from Miltenyi Biotec; and TCF-1 (S33-966) from BD Biosciences. For intracellular cytokine staining, cells were first re-stimulated in vitro with GP33-41 peptide (5 μg/mL) for 5 hrs in the presence of Golgiplug (1 μL/10^6^ cells), then fixed and permeabilized using the Cytofix/Cytoperm Kit (BD). For flow cytometry sorting, antibody-stained cells were maintained in cell sorting buffer (PBS supplemented with 1% FBS, 1% penn/strep, EDTA [2mM]) and sorted using FACSAria3 (BD), FACSAriaFusion (BD), or MoFlo Astrios EQS (Beckman). For flow cytometry analysis, antibody-stained cells were run on LSR-Fortessa X-20 analyzer (BD). All flow cytometry data were analyzed using FlowJo v.10.7.1 software (TreeStar).

### Virus titration

Viral titers were determined either by LCMV focus-forming assay as described^1^ or qPCR assay as described^2,3^.

### In Vivo cell proliferation assay

At 8 dpi., infected mice were i.p. injected with 2 μg of EdU (Beyotime, China) 4 hrs before analysis. To measure cell proliferation, cells were intracellularly stained with anti-EdU antibody and analyzed by flow cytometry. EdU staining was performed using an EdU kit (BeyoClick EdU Cell Proliferation Kit, Beyotime, China) following the manufacturer’s instructions.

### In vitro cytotoxicity assay

To generate effector T cells, naïve P14 (CD45.2^+^) cells were purified and co-cultured with a DC2.4 cell line engineered to express LCMV GP33-41 epitope. After 3 days of co-culture, P14 cells were collected. One part of splenocytes (CD45.1^+^) were incubated with 0.25 μM of CTV and pulsed with GP33-41 peptide for 1 hr and used as target cells. Another part of the same splenocytes were incubated with 5 μM of CTV but without peptide. Both parts of splenocytes were mixed in a 1:1 ratio and further co-cultured with the prepared P14 effector cells for 3 or 12 hrs at 37 °C. Survived target cells were determined by flow cytometry by comparing the percentages of peptide-pulsed vs non-pulsed splenocytes.

### In vivo cytotoxicity assay

To prepare target cells, one part of splenocytes were incubated with 0.25 μM of CFSE and pulsed with GP33-41 peptide. Another part of the same splenocytes were incubated with 5 μM of CFSE but without peptide. Both parts of splenocytes were mixed in a 1:1 ratio and co-transferred into LCMV C13 infected mice at D8 p.i. Target cell elimination was determined by analyzing the CFSE profile.

### Imaging Mass Cytometry

Samples embedded with Tissue-Tek OCT compound (Sakura) were cut into 5 μm thick tissue sections using a cryostat and mounted onto gelatin-coated histological slides, the slides were then brought to room temperature and washed 3 times with DPBS. After washing, the sections were permeabilized using 0.5 % Triton-X 100 in DPBS for 10 min and blocked with 3 % BSA in DPBS for 45 min at room temperature. A cocktail of antibodies, diluted in DPBS with 0.5 % BSA and 0.05 % Tween, was applied overnight at 4 °C at the dilutions tested in advance. The following day, slides were sequentially washed twice in 0.1 % Triton-X 100 in DPBS for 8 min, twice in DPBS for 8 min with slow agitation, followed by incubation with Iridium-conjugated intercalator (Fluidigm; 201192A) diluted 1:100 in DPBS for 30 min at room temperature. Last, slides were rinsed with ultrapure distilled water, air dried and stored at 4℃ until usage.

### Imaging Mass Cytometry data analysis and image visualization

For this purpose, we followed a procedure described previously^4^. Briefly, for each mass channel an image was reconstructed by plotting the laser shot signals in the order they were recorded and scanned to generate a pseudo-colored intensity map of each channel, and the data were subsequently examined with MCD Viewer (V.1.0.560, Fluidigm) with manually set threshold.

### scRNA-seq

For Sample Collection and Processing. Primary murine tissues were collected from co-transfer p14 mice, with tissues isolated at 5 dpi following infection with LCMV-clone 13. A total of 4 single-cell samples were prepared, consisting of CD8^+^ T cells from lung and spleen tissues, with both wild-type (WT) and knockout (KO) groups at 5 dpi. The isolated single-cell suspensions were processed using the DNBSEQ platform, which was run on the DNBelab C4 portable single-cell system (BGI). This system facilitates high-throughput cell capture and gene expression profiling. Single-cell RNA sequencing (scRNA-seq) was performed using the DNBSEQ platform from BGI, specifically the DNBelab C4 portable single-cell system. For data processing, the Cell Ranger software (v.6.1.2) was used for pre-processing, alignment, and generating feature-barcode matrices from the raw sequencing data. Subsequent quality control excluded low-quality cells based on established criteria (<500 genes or >5% mitochondrial gene content). The data was normalized and scaled to account for differences in sequencing depth across samples. Data preprocessing was carried out using Seurat v4^5^. Initial steps included filtering for low-quality cells and genes, followed by log-normalization of gene expression counts. For dimensionality reduction, principal component analysis (PCA) was performed on the highly variable genes, and the first 20 principal components were retained for subsequent clustering analysis. For visualization, t-SNE was used to embed the data into two-dimensional space, allowing for the visualization of clustering patterns. Raw sequencing data and processed analysis results will be deposited in a public database (GEO) upon acceptance of the manuscript.

### PBMCs Preparation, Purification and Activation

EDTA anticoagulated blood was collected from healthy adults. Peripheral blood mononuclear cells (PBMCs) were isolated by Ficoll-Hypaque density gradient centrifugation. T cells were purified by magnetic bead sorting followed by stimulated with PMA (50 ng/ml, Sigma-Aldrich) and ionomycin (1 µM, Sigma-Aldrich) or 5 μg/mL anti-CD3 and 2 μg/mL CD28 for 72 h in the presence of GolgiPlug™ Protein Transport Inhibitor.

### Statistical analysis

All summarized data are shown in graphs with mean ± SD. Unpaired and paired Student’s t-test were used to determine the P values. P > 0.05 was considered not significant (NS); * P < 0.05; ** P < 0.01; *** P < 0.001, **** P < 0.0001was considered statistically significant. In text and figures, “n” indicates the number of samples. All the statistical analysis was done using GraphPad Prism 8 program (GraphPad Software, San Diego, CA, USA).

## Data availability

All data are available in the main text or the extended materials. Sequence data that support the findings of this study have been deposited in the Gene Expression Omnibus (GEO) with the accession code GSE285487.

## Acknowledgments

This work was supported by the National Key R&D Program of China 2023YFC2306400, National Natural Science Foundation of China 32370956 and 32070887 (G.F.); National Natural Science Foundation of China 82273037 (H.-R. W.); National Natural Science Foundation of China 82270417 (G.L.); National Natural Science Foundation of China 32070761 (T.L.Z.); Singapore Ministry of Education MOE-000112, T2EP30220-0015 (N.R.J.G.), and Singapore National Research Foundation NRF-CRP19-2017-04 (N.R.J.G.). We also thank Dr. Haopeng Wang for kindly providing JX17 Jurkat cells.

## Author contributions

G.F., H.-R. W., E.H., G.L., B.X., N.R.J.G., J.T. and X.L.C. designed experiments. J.T., J.C.D., X.J., Q.F.Z., J.Y.W., Y.C.L., G.F.F., and X.L.C. performed experiments. J.T. and G.F. analyzed experiments. Y.Z.B. analyzed scRNA-seq data. X.J. and L.Z. performed and analyzed imaging mass cytometry with help from Y.Z.Z, S.J., J.Y.W., and others. T.L.Z., Y.Y.H., X.J., J.T. and X.Z.X. performed biochemistry experiments. B.W.H., W.Y.L., Y.L.C., T.R., Y.C., M.X.Q., C.N.Y., S.L., S.Y.L., C.H.H., L.C.H., J.B., N.G., J.R.T., C.C.X., N.W. and Y.W. contributed materials. G.F. and N.R.J.G. wrote the manuscript with review and editing from H.-R. W., E.H., G.L., B.X. Other authors contributed to reading the manuscript and providing feedback.

## Extended Data

The Extended Data includes Extended Data Fig. 1 to 9

**Extended Data Fig. 1.**
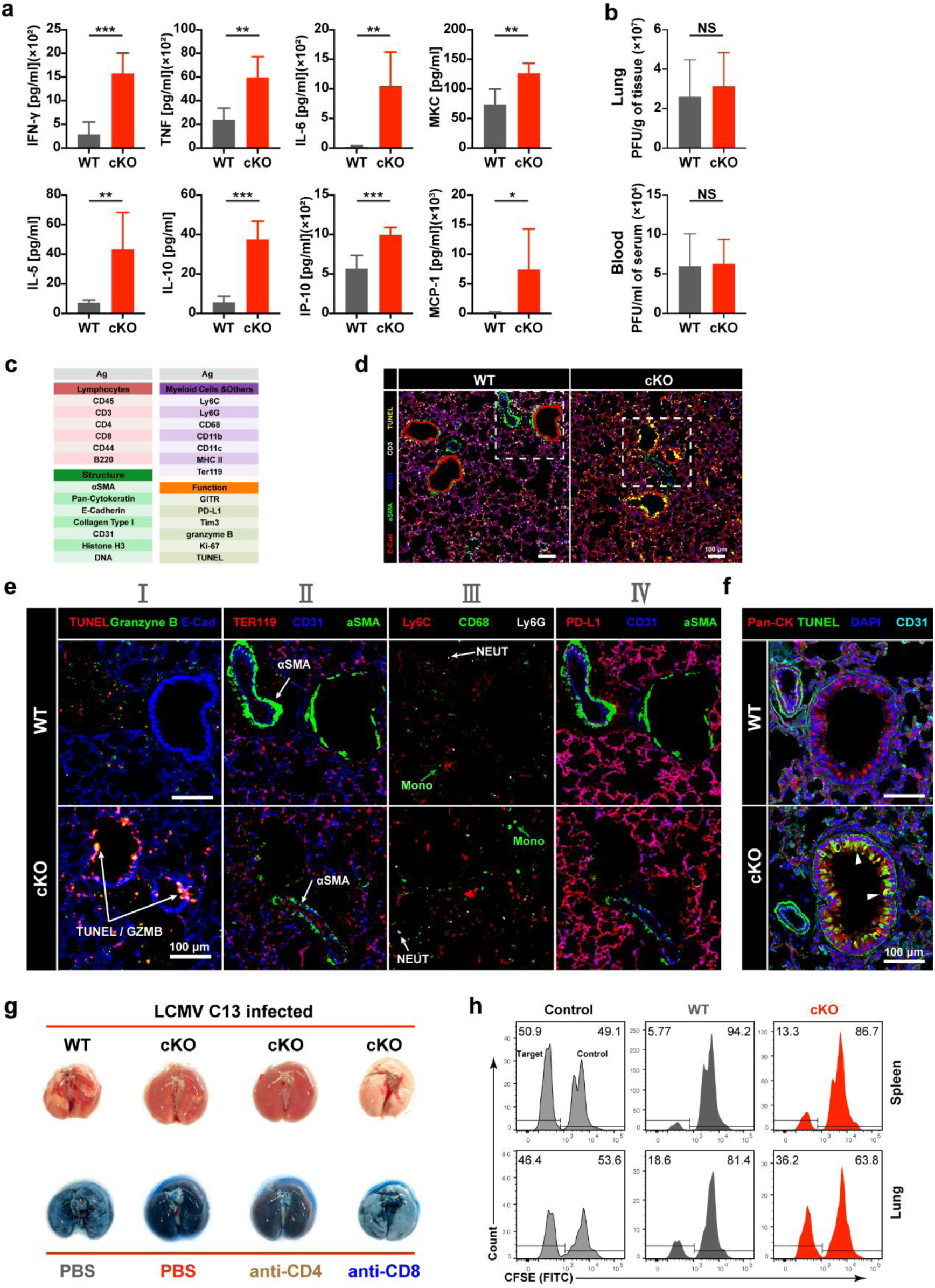
Analyses of LCMV C13-infected mice. **a**, Cytokine measurements in the BAL of mice at 8 dpi. WT (n = 6), cKO (n = 6). Data shown (mean ±SD) were from one of two experiments. **b**, Virus titers determined by plague assay at 8 dpi. Data shown (mean ±SD) were pooled from two experiments. WT (n = 16), cKO (n = 16). **c**, Antibodies used in Imaging Mass Cytometry (IMC). **d**, IMC at 1 μm resolution of lung sections. Shown is a representative 1 μm^2^ landscape view images of WT and cKO lungs respectively. **e**, IMC zoom-in images of the highlighted areas marked in (d) with indicated combinations of markers (panel I-IV) as shown in Fig. 1b. Arrows represent marker staining (I and II) or cell types (III). Scale bars, 100 μm. **f**, Confocal microscopy of lung sections in bronchi area. Arrowheads marking cell death as indicated by TUNEL staining. Scale bars, 100 μm. **g**, Lung damage of mice were evaluated by visualization (top) and Evans Blue extravasation (bottom). **h**, In vivo killing assay. Equal numbers of GP33-41 peptide-loaded target cells and no peptide-loaded control cells were labeled with 0.25 μM and 5 μM CFSE, respectively, and co-transferred into LCMV C13-infected mice at 8 dpi. Target cell elimination was determined by analyzing the CFSE profile. Data shown are from one of two experiments. Error bars show SD. Statistical comparison of experimental groups was performed using unpaired Student’s t-test (a, b). (NS, non-significant, * p < 0.05, ** p < 0.01, *** p <0.001, **** p <0.0001).

**Extended Data Fig. 2.**
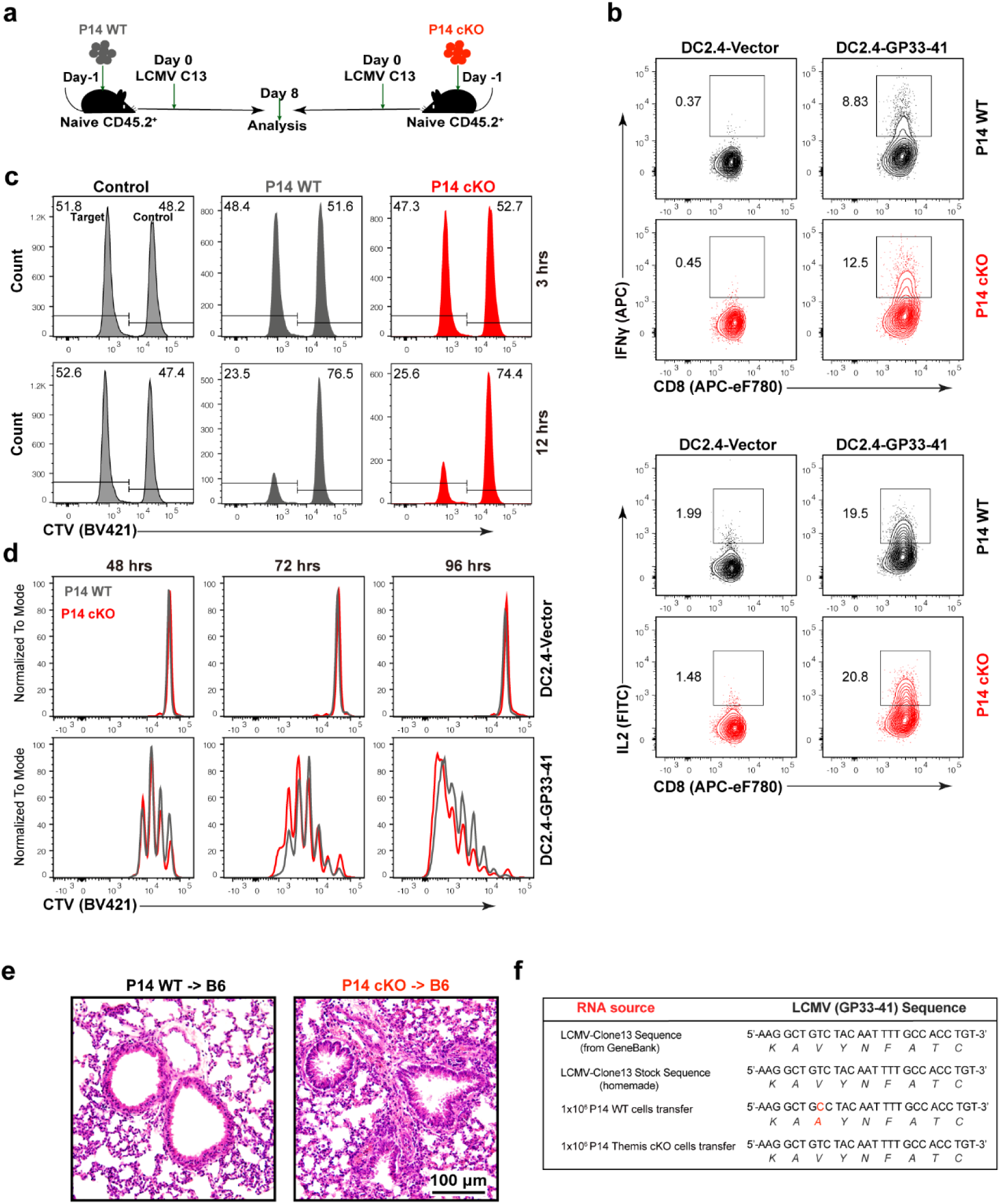
Evaluation of P14 cKO cells in vitro and in vivo. **a**, Scheme of P14 cell separate transfer experiment. **b**, In vitro cytokine assay. P14 cells were stimulated by DC2.4-GP33 cell line, and IL-2 and IFNγ expression were determined by intracellular staining. **c**, In vitro killing assay. P14 cells were stimulated by DC2.4-GP33 cell line and used as effector cells. GP33-41 peptide loaded target splenocytes and no peptide loaded control splenocytes were labeled with 0.25 μM and 5 μM of CTV respectively, mixed in a 1:1 ratio, and co-cultured with equal numbers of P14 WT or P14 cKO effector cells. Target cell elimination was determined by analyzing the CTV profile at indicated time point. Data shown were from one of two experiments. **d**, In vitro proliferation assay. CTV labelled P14 cells were co-cultured with DC2.4-GP33 cell line. CTV dilution profiles were overlaid for comparison. DC2.4-GP33 cell line express GP33-41 peptide sequence, DC2.4-vector cell line used as empty expression control. **e**, H&E staining of lung sections of infected mice receiving 1×10^6^ cells at 8 dpi. Shown are representative images. Scale bar, 100 μm. **f**, Detection of virus mutation. LCMV C13 RNA was purified from indicated source and sequenced after Q-PCR. Sequences of LCMV GP33-41 epitope were aligned for comparison.

**Extended Data Fig. 3.**
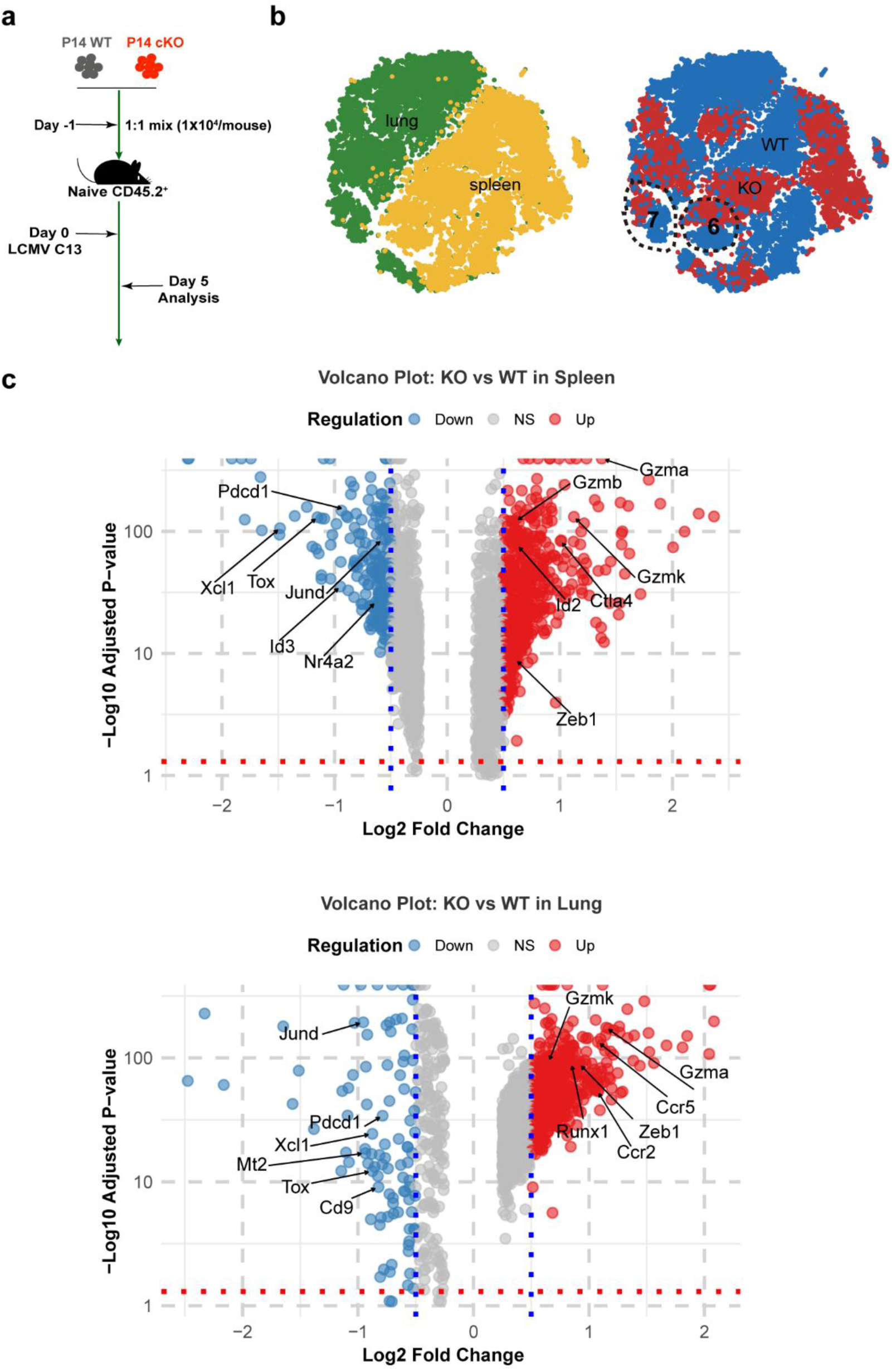
Differential gene expression analysis of co-transferred P14 cells. **a**, Scheme of P14 cell co-transfer experiment. Naive mice were co-transferred with equal numbers of P14 cells followed by LCMV C13 infection, and analyzed at indicated time points. **b**, Cells of scRNA-seq classified by organ origin (left) or genotype (right). **c**, Volcano plots illustrate the distribution of differentially expressed genes (DEGs) between cKO and WT type of P14 cells in the spleen (top) and lung (bottom) subpopulations. Genes significantly upregulated in cKO cells are highlighted in red, while those downregulated are in blue (adjusted p-value < 0.05, absolute log2 fold change > 1).

**Extended Data Fig. 4.**
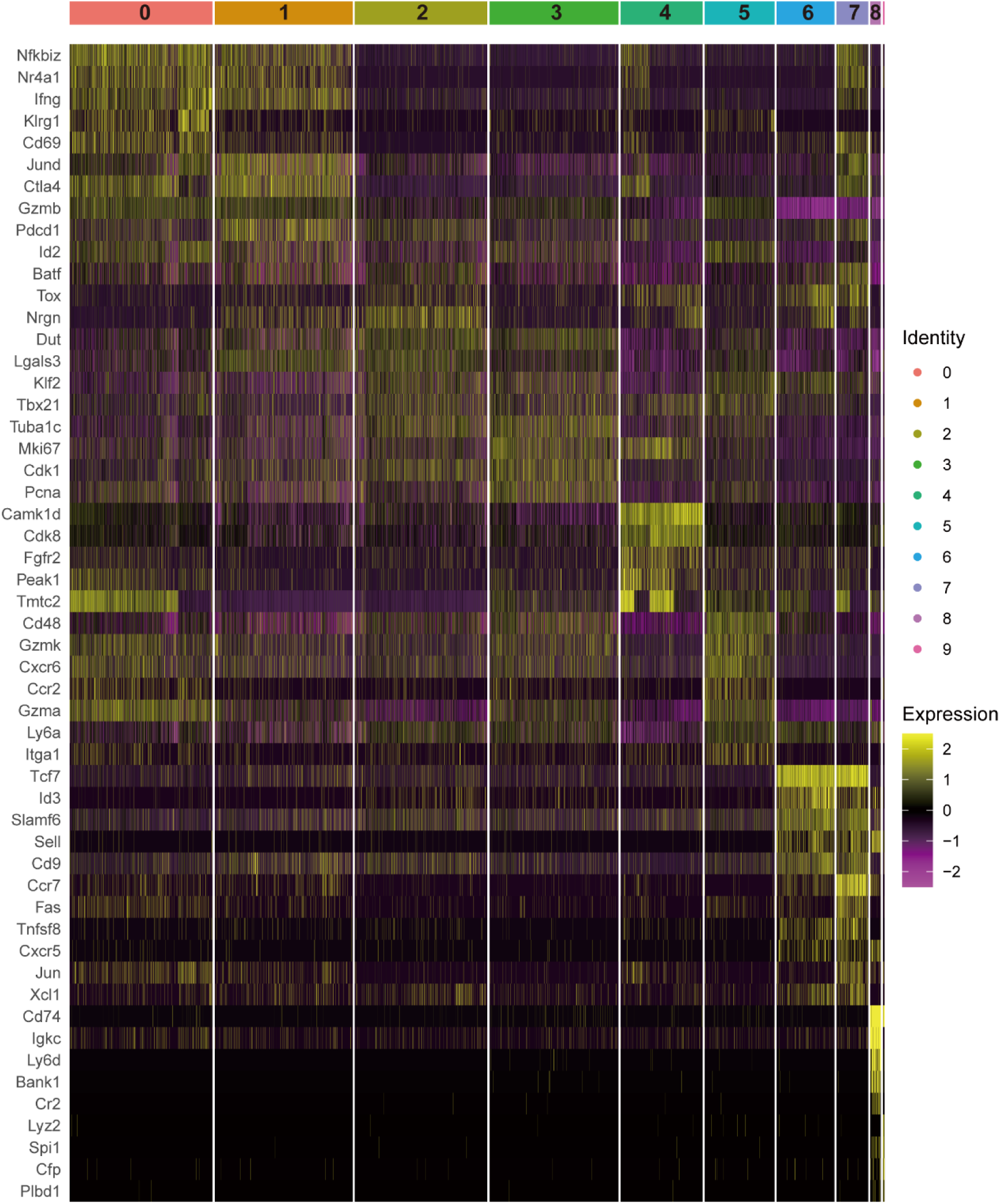
Transcriptional analysis of co-transferred P14 cells. Heatmap depicting the top 10 differentially expressed genes across individual samples, with columns representing individual cells and rows representing genes. Cells are grouped by clusters, and the color scale, based on z-scores, ranges from -2 (deep purple, indicating low expression) to +2 (vivid yellow, denoting high expression).

**Extended Data Fig. 5.**
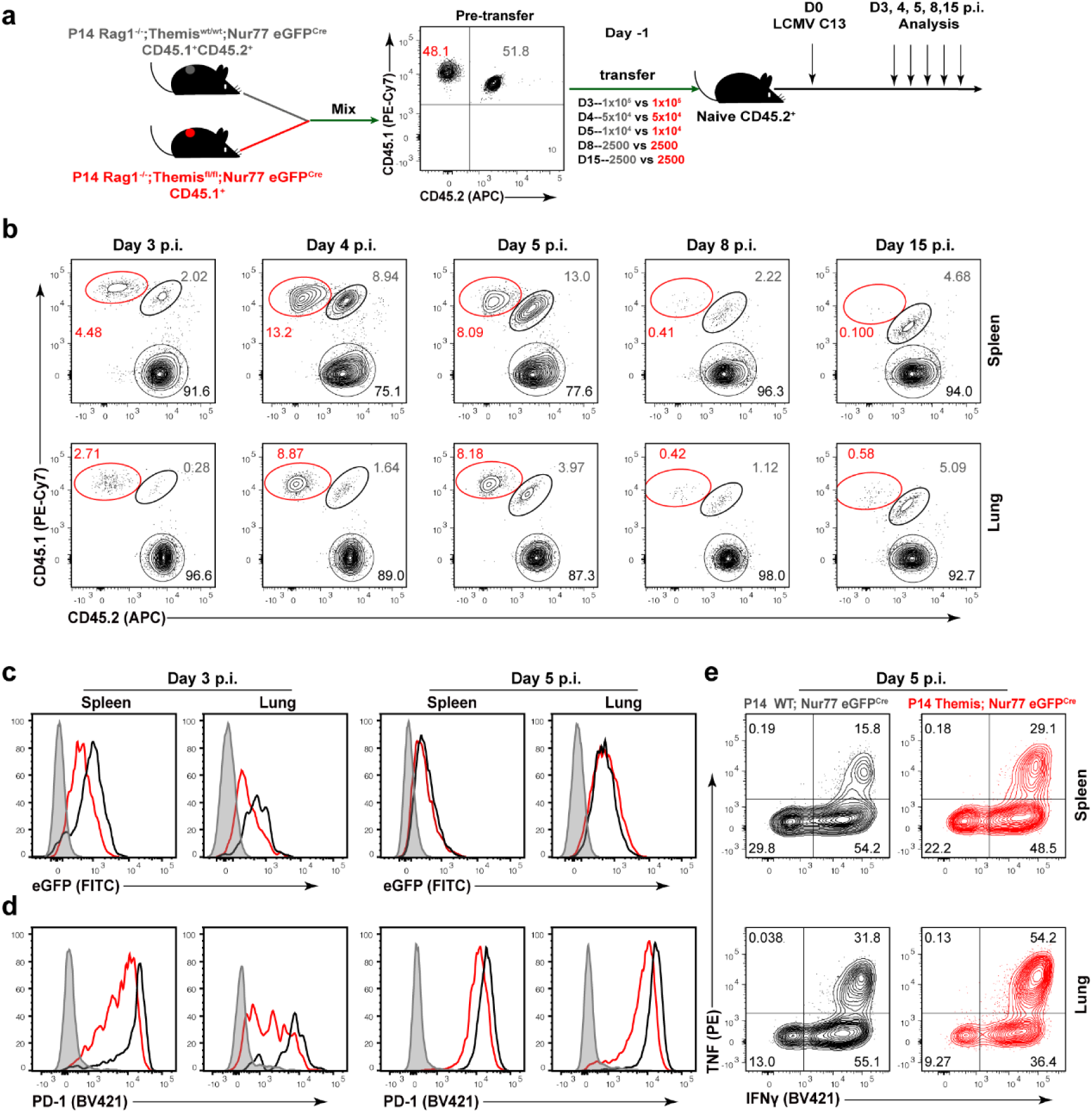
Orderly regulation of TCR and PD-1 signaling by Themis. **a**, Scheme of experimental setup. Naive mice were co-transferred with equal numbers of P14 WT/Nur77^GFP^ and P14 cKO/Nur77^GFP^ cells followed by LCMV C13 infection and analyzed at indicated timepoint. **b**, Representative FACS plots showing the proportions of P14 WT/Nur77^GFP^ and P14 cKO/Nur77^GFP^ cells at indicated time points. **c**,**d**, Representative FACS plots showing GFP (c) and PD-1 (d) expression in P14 cells at indicated time points. **e**, Representative FACS plots showing the expression of IFNγ and TNF in P14 cells at 5 dpi. Data shown are from one of two experiments.

**Extended Data Fig. 6.**
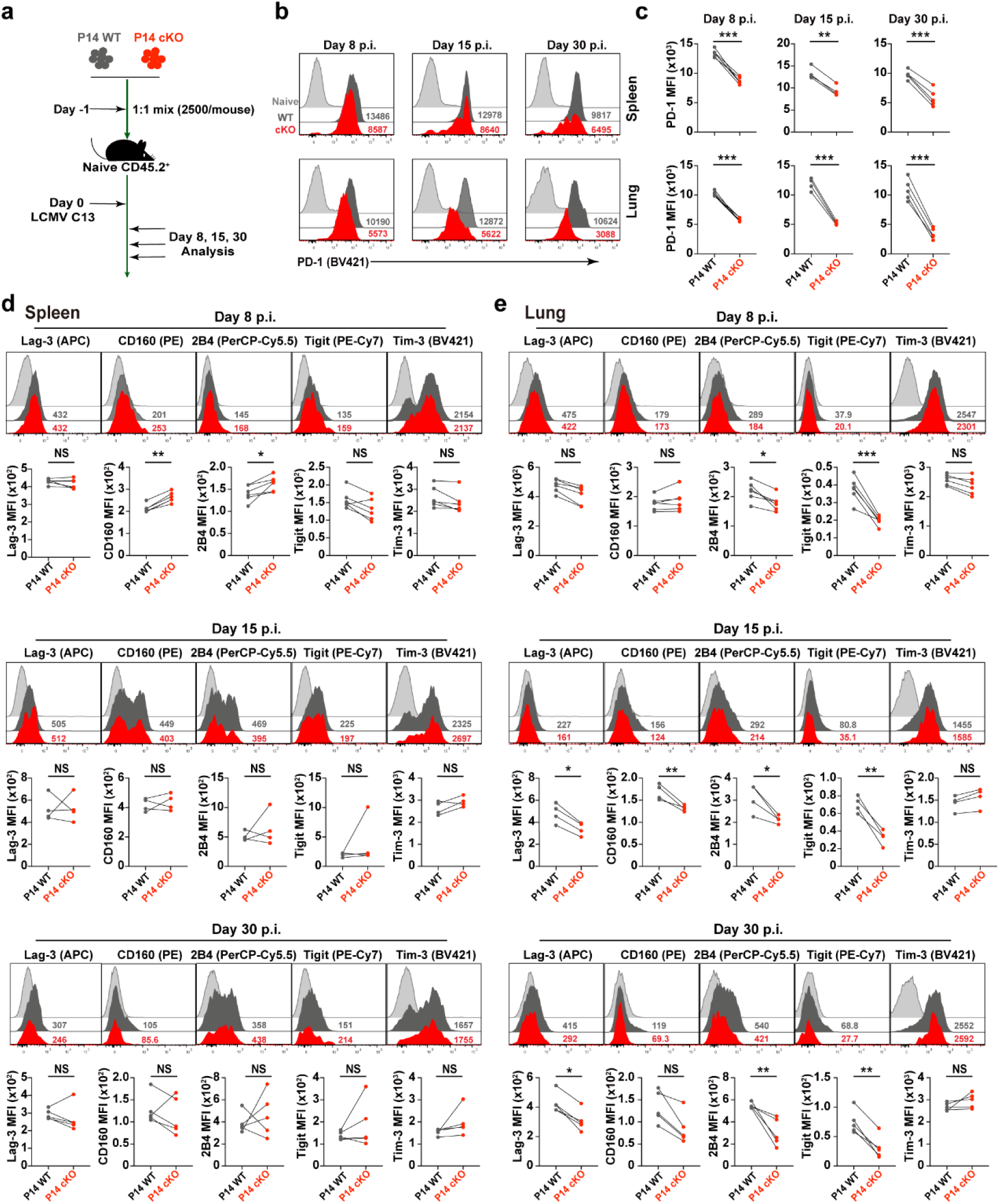
Expression of PD-1 and other co-inhibitory molecules. **a**, Scheme of experimental setup. Naive mice were co-transferred with equal numbers of P14 cells followed by LCMV C13 infection, and analyzed at indicated time points. **b**,**c**, Expression of PD-1 in co-transferred P14 cells in the spleen and lung. Shown are representative histograms at the indicated time point (b) and summary of data (c). **d**,**e**, Expression of indicated co-inhibitory molecules in co-transferred P14 cells in the spleen (d) and lung (e). Shown are representative histograms at indicated time point (top row) and summary of data (bottom row). MFI, mean fluorescence intensity. Experiments were performed three times (b, c) and two times (d, e) with ≥ 4 mice per timepoint of each group. Statistical comparison of experimental groups was performed using paired Student’s t-test. Error bars show SD. (NS, non-significant, * p < 0.05, ** p < 0.01, *** p <0.001, **** p <0.0001).

**Extended Data Fig. 7.**
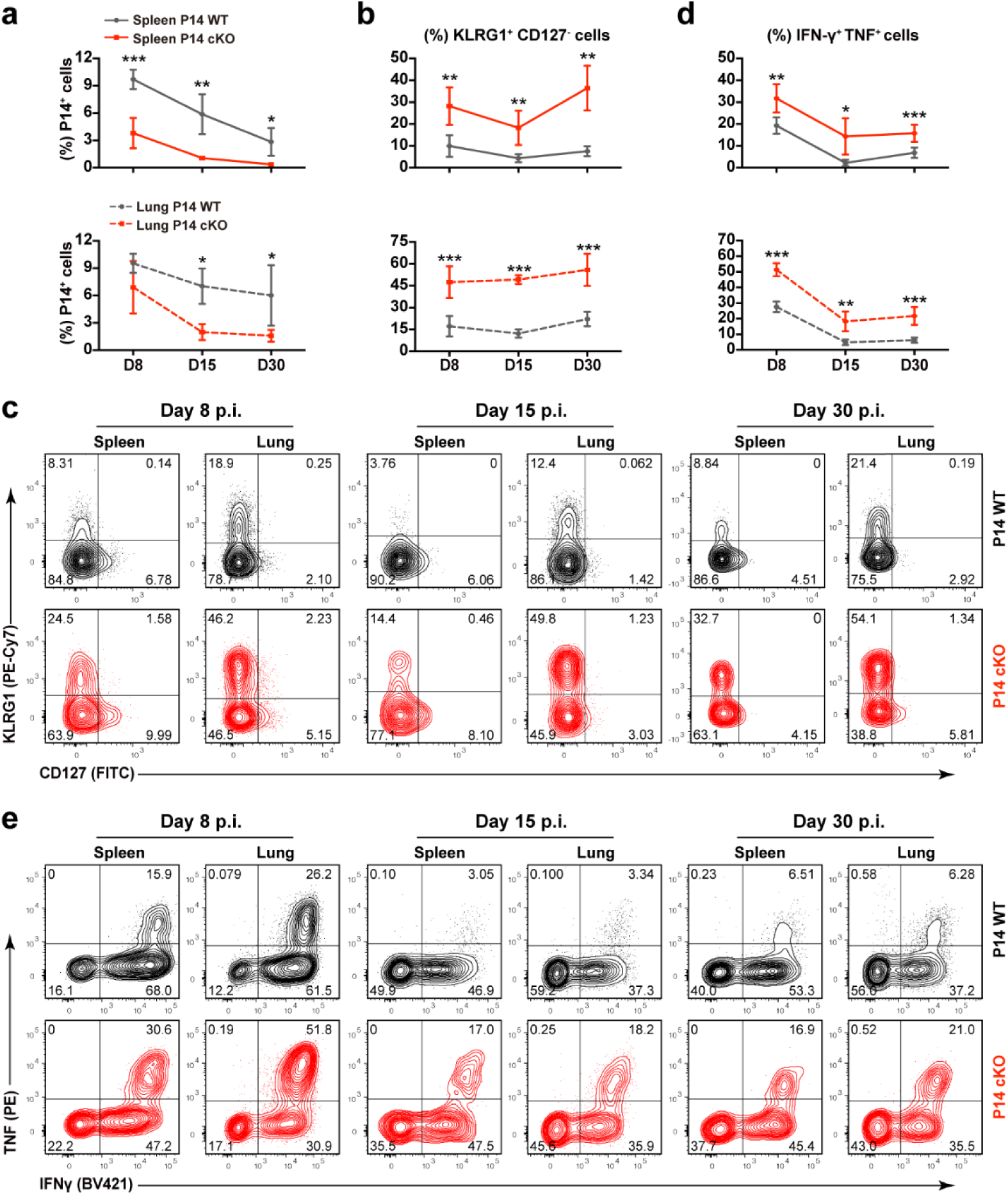
Themis is required for the maintenance of T-ex cells. **a**, The kinetic changes of the proportion of P14 cells. **b**,**c**, Expression of KLRG1 and CD127 in co-transferred P14 cells at indicated time points. Shown are summary of data (b) and representative FACS plots (c). **d**,**e**, Expression of IFNγ and TNF in co-transferred P14 cells at indicated time points. Shown are summary of data (d) and representative FACS plots (e). Experiments were performed two times with ≥ 4 mice per timepoint of each group. Statistical comparison of experimental groups was performed using paired Student’s t-test. Error bars show SD. (NS, non-significant, * p < 0.05, ** p < 0.01, *** p <0.001, **** p <0.0001).

**Extended Data Fig. 8.**
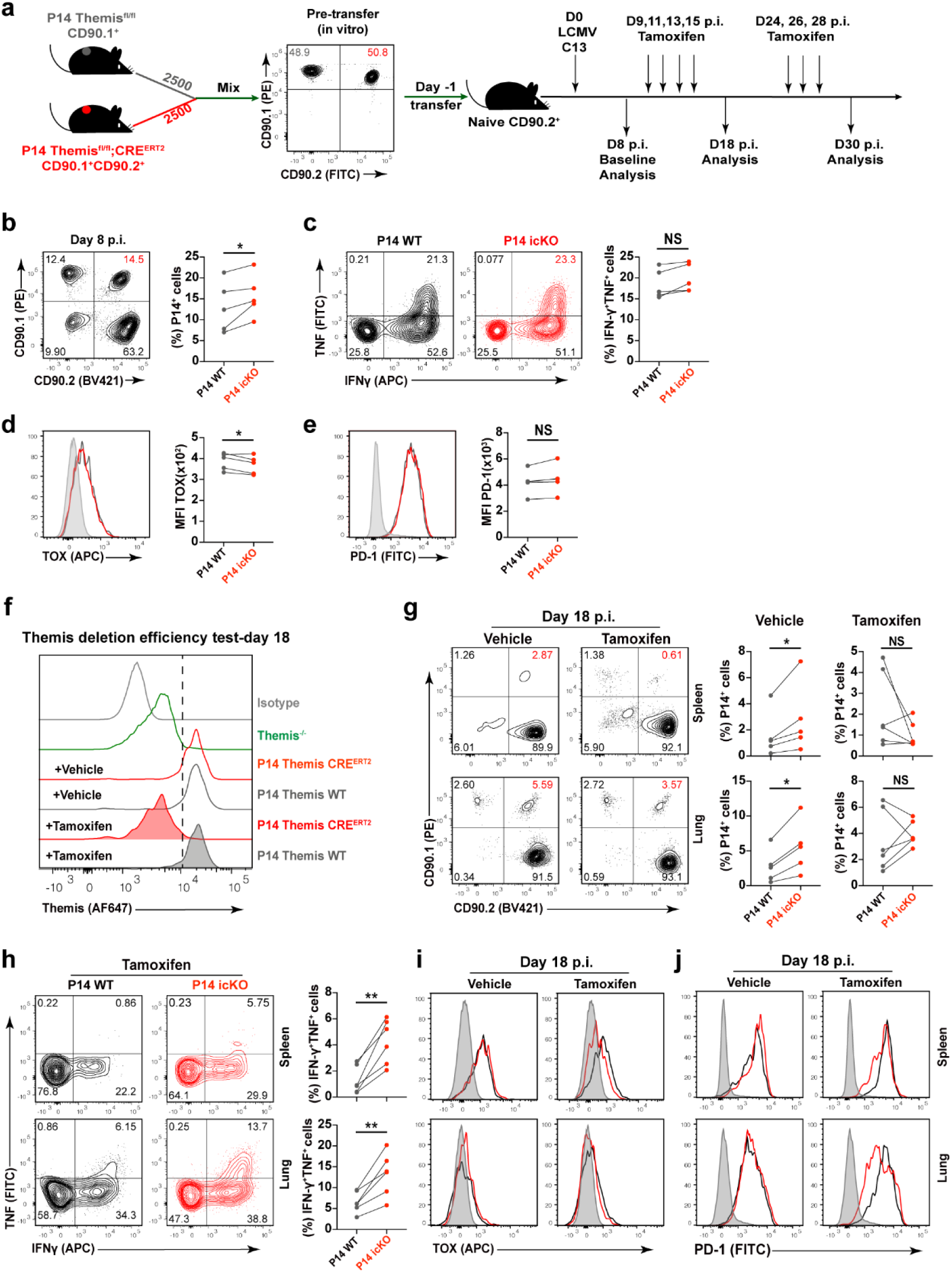
Baseline analysis of P14 icKO cells. **a**, Scheme of experimental setup. P14 Themis^flox/flox^ Cre^ERT2^ mice derived cells (P14-icKO, CD90.1^+^/ CD90.2^+^) and P14 Themis^flox/flox^ mice derived cells (P14 WT, CD90.1^+^) were co-transferred into naive CD45.2^+^ recipients followed by LCMV C13 infection. Tamoxifen treatment was applied at indicated time points. **b**-**e**, Baseline analyses at 8 dpi before tamoxifen treatment. Shown are representative FACS plots and summary of data for the proportion of P14 cells (b), the expression of IFNγ and TNF (c), TOX (d), and PD-1 (e). **f**-**j**, Analysis of P14 cells at 18 dpi after one round of tamoxifen treatment. (f) Themis deletion verification by intracellular staining. Shown are histograms of indicated condition. (g) The proportion of P14 cells after vehicle or tamoxifen treatment. Shown are representative FACS plots and summary of data. (h) Expression of IFNγ and TNF in co-transferred P14 cells after tamoxifen treatment. Shown are representative FACS plots and summary of data. i,j, Expression of TOX (i) and PD-1 (j) in co-transferred P14 cells at indicated time points after vehicle or tamoxifen treatment. Shown are representative histogram overlays of P14 WT and P14 icKO cells. Gray shadows represent splenocytes from naive mouse, used as control. Experiments were performed two times per timepoint of each group. Statistical comparison of experimental groups was performed using paired Student’s t-test. Error bars show SD. (NS, non-significant, * p < 0.05, ** p < 0.01, *** p <0.001, **** p <0.0001).

**Extended Data Fig. 9.**
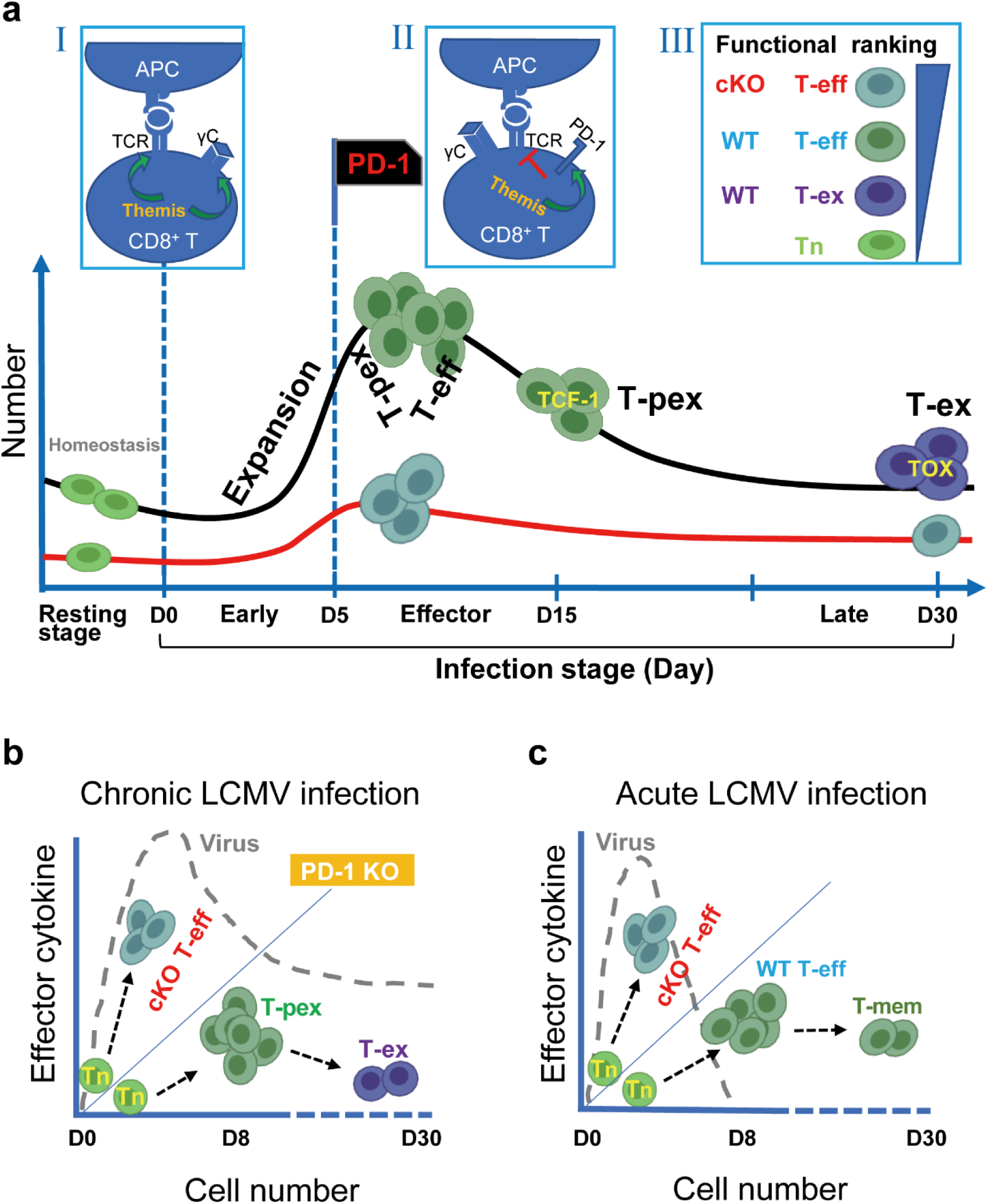
Themis orderly regulates TCR and PD-1 signals. **a**, Comparison of virus-specific CD8^+^ T cells responses in WT and Themis-cKO mice. Themis-deficient CD8^+^ T cells have a “reduced number” feature at the resting stage, partially due to their impaired responsiveness to common γc cytokines (Box I). This trend continues during the early infection stage when naive (Tn) cells expand and differentiate into short-lived T-eff cells. During the effector stage, PD-1-mediated inhibition begins to suppress the function of WT T-eff cells, but fails to do so in cKO T-eff cells, either due to its impaired expression or defective signaling, or both (Box II). Therefore, cKO T-eff cells are more functional than WT T-eff cells due to the “enhanced function” feature (Box III). Under normal circumstances, some WT CD8^+^ T cells differentiate into TCF-1^+^ T-pex cells and finally differentiate into TOX^+^ T-ex cells, however, the development of both cell types is severely blocked in cKO CD8^+^ T cells. **b**, In chronic LCMV infection, WT CD8^+^ T cells maintain a balance between cell number and function and undergo normal T cell exhaustion. In contrast, cKO CD8^+^ T cells have reduced numbers but enhanced function, whereas PD-1 KO CD8^+^ T cells have both increased numbers and functions. Importantly, in the latter two cases, T cell exhaustion was interrupted and Themis cKO and PD-1 KO mice died during chronic LCMV infection. **c**, In acute LCMV infection, although the intrinsic properties of cKO CD8^+^ T cells remain the same as in chronic infection, mice survive because the virus is cleared.

